# A pan-cancer landscape of somatic substitutions in non-unique regions of the human genome

**DOI:** 10.1101/2020.04.14.040634

**Authors:** Maxime Tarabichi, Jonas Demeulemeester, Annelien Verfaillie, Adrienne M. Flanagan, Peter Van Loo, Tomasz Konopka

**Affiliations:** Francis Crick Institute, London, UK; KU Leuven, Department of Human Genetics, Leuven, Belgium; Research Department of Pathology, Cancer Institute, University College London, London, UK; Department of Cellular and Molecular Pathology, Royal National Orthopaedic Hospital NHS Trust, Stanmore, Middlesex, UK; William Harvey Research Institute, Queen Mary University of London, London, UK

**Author notes:** Equal contribution (Peter Van Loo, Tomasz Konopka).

## Abstract

Around 13% of the human genome displays high sequence similarity with at least one other chromosomal position and thereby poses challenges for computational analyses such as detection of somatic events in cancer. We here extract features of sequencing data from across non-unique regions and employ a machine learning pipeline to describe a landscape of somatic substitutions in 2,658 cancers from the PCAWG cohort. We show mutations in non-unique regions are consistent with mutations in unique regions in terms of mutation load and substitution profiles, and can be validated with linked-read sequencing. This uncovers hidden mutations in ~1,700 coding sequences and thousands of regulatory elements, including known cancer genes, immunoglobulins, and highly mutated gene families.

## Introduction

Catalogs of somatic mutations in cancer promise insights into disease-initiating pathways, underlying evolutionary processes, and the identification of potential therapeutic opportunities. Large-cohort studies, for example The Cancer Genome Atlas (TCGA, https://www.cancer.gov/tcga) and the International Cancer Genome Consortium (ICGC, https://dcc.icgc.org/), have shed light on the complexity of the mutational landscapes in gene-coding regions. The Pan-Cancer Analysis of Whole Genomes (PCAWG) study has further undertaken the analysis of 2,658 whole cancer genomes from the ICGC and TCGA to characterize regions unobserved via exome sequencing studies^1^. This resource has led to studies of the processes underlying somatic events^2^, complex structural variation^3^, driver genes^4^, and timing^5^. However, these analyses rely on variants that can be positioned uniquely in the genome and thus the repertoire of variation in non-unique regions still remains unexplored.

Short-read sequencing – the technology used in cancer studies including PCAWG – identifies somatic mutations by comparing fragments of DNA from normal and tumor samples to a reference genome. At the scale of 100bp, however, 13% of the human genome consists of sequences that are present at more than one chromosomal location^6,7^. These regions range in multiplicity from two to several thousand copies and in identity of sequences from vague to perfect matches. This non-uniqueness complicates genetic analyses and creates recurrent blind spots to somatic mutation calling. Irrespective of their amenability to analysis, non-unique regions include genes and regulatory elements that participate in human diseases^8^, developmental processes^9^, as well as splicing factors and nuclear RNAs that are recurrently mutated in cancers^10,11^. Blind spots in these regions thus hinder a systematic understanding of relevant biological processes.

To alleviate the limitations of variant detection from short-read sequencing due to non-unique regions, a technique called thesaurus annotation characterizes mutations in terms of equivalence classes of genetic positions^12,13^. This approach does not pinpoint the precise location of mutations, but it enables calculation of summary statistics such as mutational load. It also provides sufficient information about somatic events to study mutational signatures^14^ and to identify affected functional elements. In the present work, we employed this technique to study the PCAWG dataset with the aim to describe the landscape of somatic single-base substitutions. This uncovers a vast set of somatic events across cancer genomes and cancer types.

## Results

### Thesaurus annotation uncovers a distinct class of mutations in a pan-cancer cohort

To perform an analysis of somatic mutations that is inclusive to non-unique regions, we set up a pipeline for variant calling on the PCAWG dataset without filtering reads based on mapping quality (Methods). This provided a comprehensive set of candidate positions in all regions of the genome. We then annotated the sites using a procedure that links called sites to possible alternative positions in the genome^12^ (Figure 1a). To search for somatic mutations among the candidates, we utilized curated PCAWG data in two distinct ways. First, we assembled a panel of 237 genomes from normal tissues to filter out common germline polymorphisms and sequencing artifacts. Second, we trained a machine learning model to classify somatic events. Similar to other machine-driven approaches^15,16^, our pipeline provided an algorithm with 18 features about candidate sites collected from tumors and matched normal samples. The algorithm then learned a strategy to call mutations from the candidates to match the PCAWG consensus calls^1^. Crucially, training and testing were performed using only data from unique regions where the PCAWG call set is expected to be of high quality, and the features provided did not include information about mapping quality. The final classifier achieved a root-mean-square error of 8.9%, with false discovery and false negative rates of 7.6% and 4.3%, respectively (Figure S1). The most important feature for classification was allelic frequency in the matched normal sample (increase in classification error to 37% when omitted), but many other features, such as allelic frequency in the tumor and the frequency in the panel of normals contributed as well (Figure S2).

**Figure 1.**
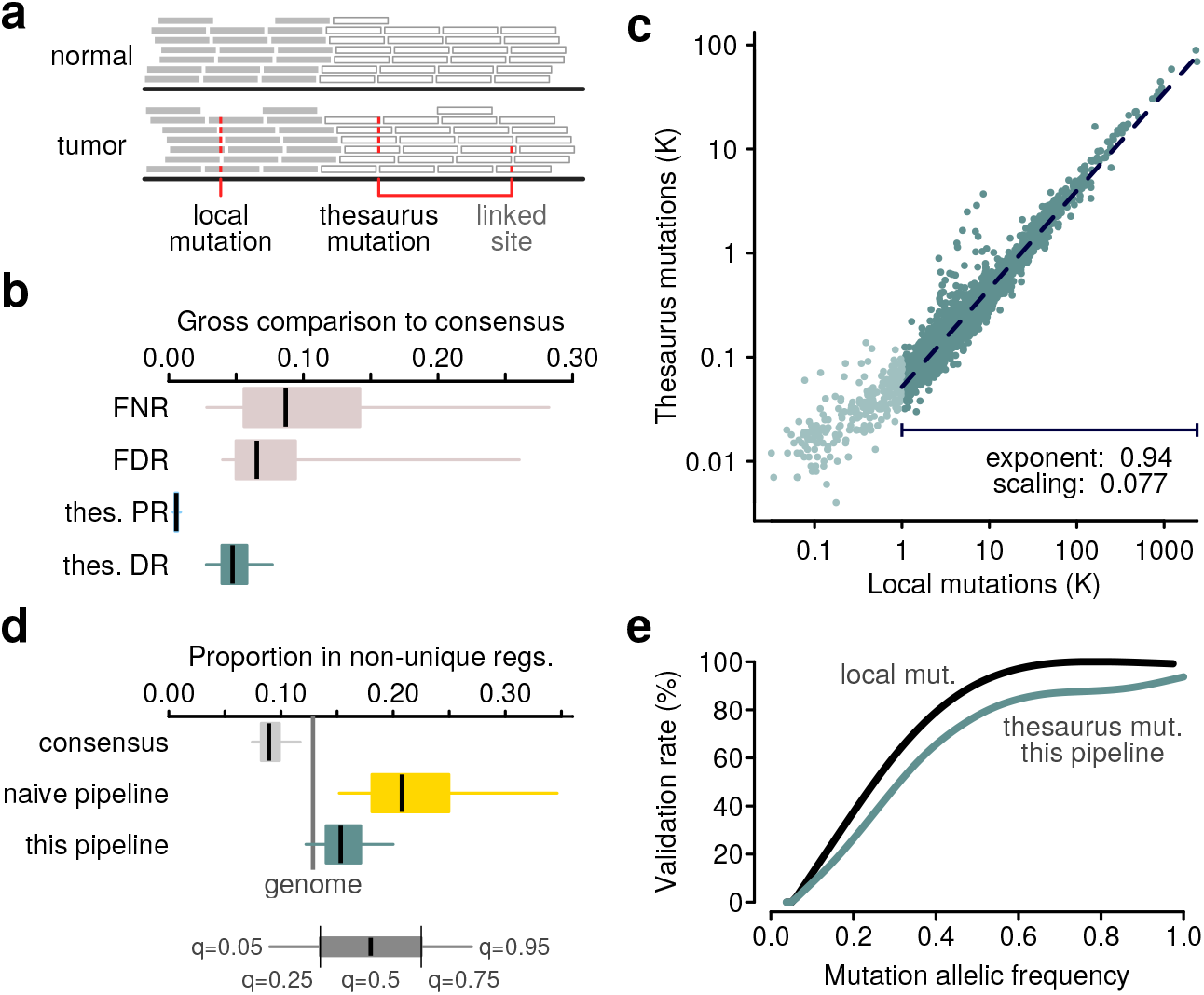
Calling mutations in non-unique regions of the genome. **(a)** Schematic of short-read sequencing data with one single-nucleotide substitution in a unique region and one in a non-unique region. Thesaurus annotation links related sites together. **(b)** Comparison of mutation calls from a mappability agnostic pipeline with the PCAWG mutation set. FDR: false discovery rate; FNR: false negative rate; thes. PR: positive rate among thesaurus mutations; thes. DR: new discovery rate among thesaurus mutations. **(c)** Mutation load among simple and thesaurus mutations. A model is fit on a subset of samples with at least 1000 simple mutations. **(d)** Proportions of mutations that fall in non-unique regions of the genome. Vertical bar shows the fraction of the genome that is non-unique at a resolution of 100bp. The cohort was filtered to exclude samples with fewer than 1000 simple mutations. consensus: PCAWG mutation calls; naive: a mutation set called considering non-unique regions, but without thesaurus annotations. **(e)** Validation rate of mutations in an independent cancer sample sequenced with short-read and linked-read technologies.

After training the machine-learning model, we processed the entire PCAWG dataset and thereby produced new sets of somatic mutation calls. We split mutations in the new call set into those placed uniquely in the genome, which we describe further as ‘simple’ or ‘local’, and those that can be linked to alternate sites, which we term ‘thesaurus’. Compared to the PCAWG calls, our set of simple mutations showed median false discovery and false negative rates per sample at 7% and 9%, respectively, albeit with variability across the samples in the cohort (Figure 1b). Such discrepancies and variability are not unexpected, as differences among computational pipelines are well-documented^17,18^. Indeed, modeling revealed that mutation frequency, coverage, and mutation spectrum can explain the largest discrepancies, and that high false discovery and false negative rates are related to internal consistency within the consensus itself (Figure S3). Performance was stable across cancer types (Figure S4).

Importantly, the set of thesaurus mutations showed little overlap with the PCAWG calls (Figure 1 b), indicating most of those sites were previously hidden. To investigate whether these sites are reasonable additions to the samples’ landscape, we studied total mutation load across samples among the simple and thesaurus calls and found a high correlation (spearman rho 0.96) that was concordant with direct proportionality (Figure 1c). Other properties such as allele frequency and mutation coverage were also concordant (Figure S5). Moreover, counts of thesaurus mutations correlated only weakly with sequencing coverage (spearman rho 0.16), suggesting the calls were not dominated by noise.

We then studied the position of mutations in relation to annotations of non-unique sequence. As expected, PCAWG calls showed under-representation in regions with low mapping quality (Figure 1d). A simulation of mutation calls that might be obtained using a naive pipeline – considering loci in non-unique regions but without using thesaurus annotation – showed severe over-representation (Figure 1d), providing a justification for common mutation callers to filter out such regions. In contrast, our pipeline produced an intermediate distribution, albeit with a systematic over-representation compared to the proportion of non-unique sequence. This can be due to a residual level of false positives, due to gaps in the sequence of the human reference genome, or due to a propensity for samples to accumulate or tolerate mutations in those regions.

As an orthogonal validation, we performed short-read and linked-read sequencing on one additional cancer sample (Methods). The linked-read protocol uses barcodes to help aligners position reads at their correct coordinates in the reference genome^19,20^ and thus expands the regions where variation can be assessed by common mutation callers. In the short-read data, our pipeline called 3,074 simple and 189 thesaurus somatic mutations, which we sought to confirm in the linked-read data. Validation rates for simple variants were proportional to the variant allele frequency in the short reads and surpassed 90% at allele frequency of 0.5 (Figure 1 e, Figure S6). For calls with thesaurus annotation, the validation rate was just 11% lower. This suggests that while thesaurus calls may retain more false components than local calls, the large majority of hits from our pipeline represent real events.

### Mutations in non-unique regions are consistent with known mutational processes

Somatic mutations in cancers appear through several biochemical processes, many of which leave distinct patterns in samples’ mutation profiles^2,14^. To a first approximation, these processes can be presumed to act similarly in unique and non-unique regions and therefore manifest among simple as well as thesaurus-annotated mutations. To test this hypothesis, we stratified mutations by trinucleotide contexts. These profiles, which consist of 96-dimensional vectors, were correlated in most samples (Figure 2a). Investigations of representative samples (Figure 2b) suggested that the strength of correlation was influenced by the mutation load. Indeed, modeling revealed that 86% of discrepancies between simple and thesaurus profiles could be accounted through mutation load, the entropy of the trinucleotide profiles, and technical features such as depth of coverage.

**Figure 2.**
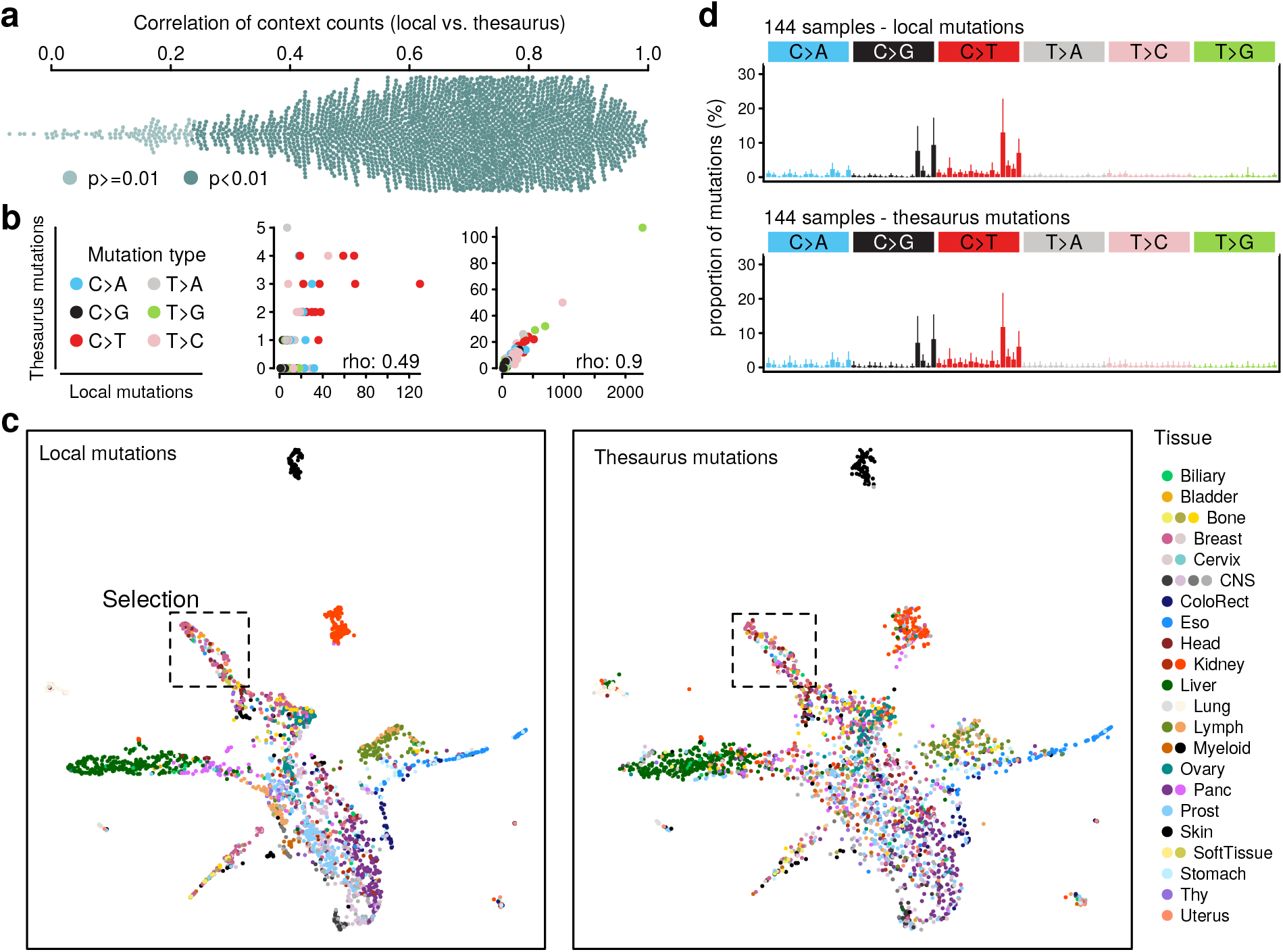
Concordance of simple and thesaurus mutational profiles. **(a)** Distribution of spearman correlations, computed for each sample, between 96-dimensional mutational profiles from simple and thesaurus mutations. **(b)** Correlations diagrams between thesaurus and simple mutation counts for two representative samples with intermediat and high correlation values. **(c)** Stereo UMAP embedding of cohort samples based on trinucleotide mutation profiles. The first map is based on simple mutations and a cosine distance. The second view shows a projection of thesaurus profiles onto the same embedding space. Colors indicate histology types and subtypes. **(d)** Mutation profiles for local and thesaurus mutations for a group of samples selected from the dashed area in (c). Mutation profiles display average profiles for the group. Bars represent 95% quantiles for each substitution type.

To summarize the heterogeneity of the mutation profiles across the cohort, we visualized the similarities between samples in a low-dimensional embedding (Figure 2c). In contrast to analyses that decompose mutation profiles into independent signatures^2,14^, this technique compares samples in a holistic manner. Profiles based on simple mutations clustered into several distinct groups, reproducing known characteristics of the pan-cancer cohort. For example, skin and kidney cancers separated from all others in this visualization, indicating that their mutation profiles are formed by distinct combinations of mutational processes. We then asked to what extent thesaurus mutations capture the same patterns and projected the thesaurus profiles onto the same visualization. Because of the lower overall counts, the resulting patterns were noisier, but nonetheless mirrored the original. This global picture was confirmed by focusing on sets of samples from distinct areas of the embedding (Figure 2d, Fig S7). Altogether, thesaurus mutations have similar characteristics as simple mutations across cancer types as well as similar molecular mutagenic processes.

### Thesaurus mutations affect thousands of functional elements

While most mutations in cancer genomes are passengers, some inflict functional effects, for example, by modifying protein structure or altering gene regulation. To create a comprehensive summary of the impact of thesaurus mutations in tumors, we partitioned the genome into nonoverlapping regions described by a gene identifier and a functional label (coding sequence, intron, promoter, untranslated, intergenic). These regions are defined through gene annotations, not sequence uniqueness, and thus we found that some carried only simple mutations, others contained only thesaurus mutations, and others harbored both types. In aggregate, thesaurus mutations were associated with thousands of genes, including 1,744 coding sequences (Figure 3a).

**Figure 3.**
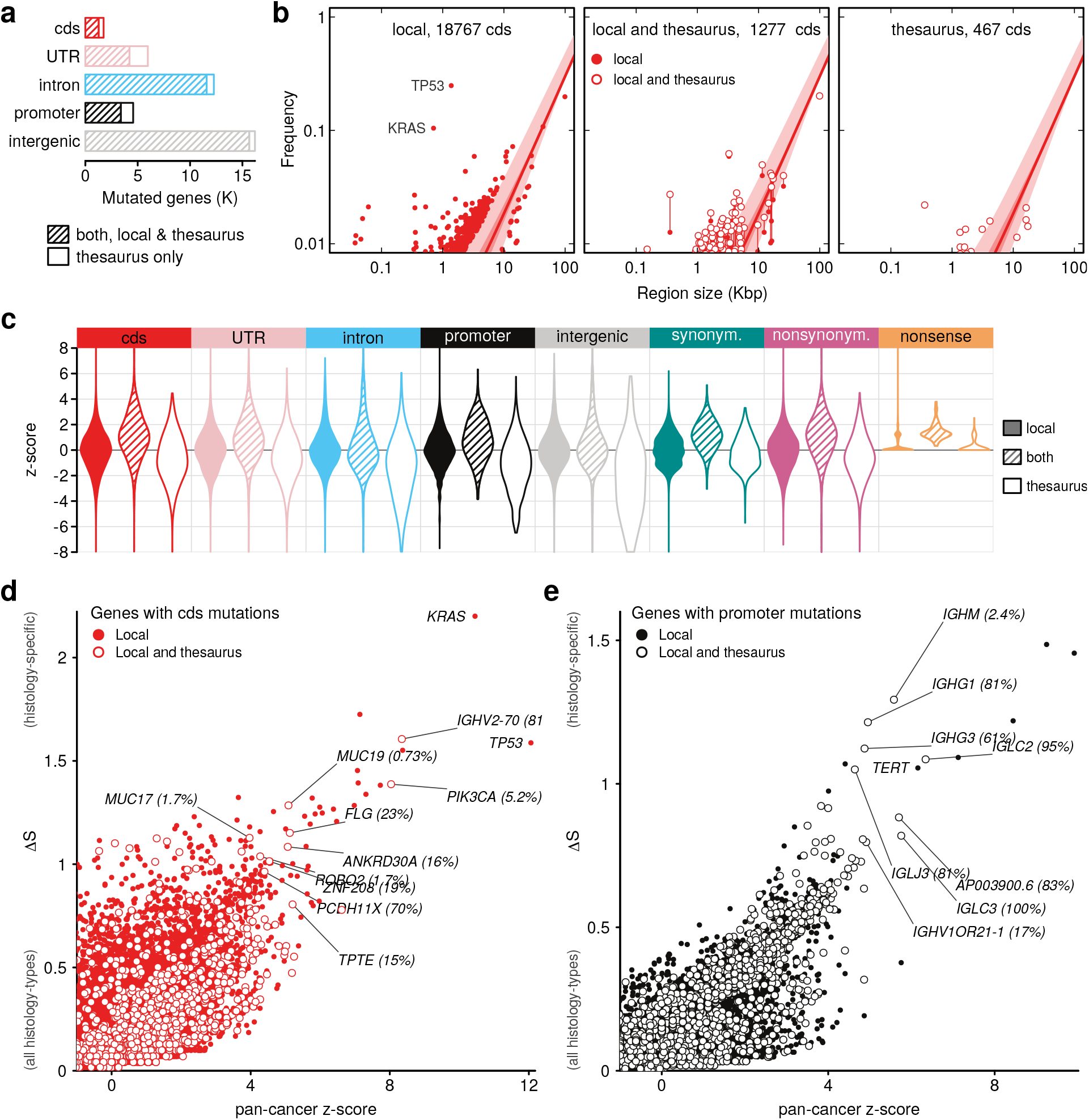
Mutation rates in functional regions. **(a)** Counts of functional regions affected by both local and thesaurus mutations, or only by thesaurus mutations. **(b)** Modeling of cohort frequency in gene coding sequences (cds) as a function of region size using quantile regression. Shaded regions indicate quantile 25% and 75% intervals and dots display genes outside the range. Subpanels show genes affected only by local mutations, by a mixture of local and thesaurus mutations, or only thesaurus mutations. **(c)** Summary of quantile modeling on functional regions, and in coding sequences partitioned by mutation effect. **(d)** Specificity of mutations in coding sequences. Two axes measure over-representation of mutations in the pan-cancer cohort and the entropy across cancer types. Labels indicate two canonical cancer genes – TP53 and KRAS – and outlier genes that contain at least some thesaurus mutations. Percentages show the proportion of samples with thesaurus mutations. **(e)** Analogous to previous panel, showing promoter regions.

Several strategies are used to assess the importance of mutations and identify driver genes^21–24^. Recurrence in a cohort is a key indicator, but this signal can be confounded by factors such as region size, chromosomal location, proximity between adjacent mutations, sequence composition, and, in the case of coding sequences, effects on protein structure^22,25^. However, these covariates can themselves be confounded by non-uniqueness in the genome. Here, in order to study mutation patterns across all region types and in a way compatible with thesaurus candidates, we performed modeling using only region length as a covariate (Methods). Starting with coding sequences, we fit quantile regression models to describe cohort frequency in genes with unique sequence (Figure 3b). The resultant model captured the expected upward trend, with established driver genes such as *TP53* and *KRAS* as strong outliers. Applying the same model on genes that include non-unique sequences revealed the same trends. Interestingly, the cohort frequencies of many genes shifted across quantile boundaries depending on whether thesaurus mutations were excluded or included.

As the same modeling strategy is also suitable to study mutations outside of coding regions, we carried out a genome-wide analysis and summarized the deviation of individual regions from model trends using z-scores (Figure 3c, Table S1). Distributions of z-scores for elements affected only by local mutations centered around zero by construction. For regions affected by thesaurus mutations, distributions were also centered near zero despite this property not being built into the models. There was a consistent shift toward positive z-scores, but distributions for genes harboring thesaurus mutations exclusively included heavy tails of negative scores, indicating the mutation set may still suffer from false negatives. Scores were consistent when the same modeling was repeated on sub-cohorts and, for coding regions, were correlated with a ranking produced by a specialized scheme accounting for additional covariates (Methods, Figure S8). Overall, the z-scores therefore provide a reasonable, albeit rudimentary, prioritization of hits.

To further refine the prioritization, we computed an entropy-based measure of specificity across cancer types (Methods). We then used the pan-cancer z-scores and specificity measures together to visualize hits in coding sequence (Figure 3d), promoters (defined as regions of at most 2000bp upstream of genes, Figure 3e), and other regions (Figure S9, S10, Table S1). This approach captured expected characteristics, in particular that most genes are neither recurrently mutated or specific to a cancer-type, and that both dimensions carry outliers. Coding regions in *TP53* and *KRAS* were the top hits for pan-cancer recurrence and specificity, respectively. Genes with thesaurus mutations lay in intermediate regions of the distributions interspersed among other cancer genes. Strikingly, top-ranked thesaurus genes included well-known cancer genes such as *PIK3CA,* including thesaurus mutations in breast cancers. Another pattern, visible in the analysis of coding sequences, but that was even more pronounced among promoter sequences (Figure 3e), was high recurrence and specificity among immunoglobulin elements of the *IGLC, IGHG, IGHJ, and IGHM* families. These instances offer leads for more in-depth investigation of thesaurus hits in selected gene families.

### Thesaurus annotation uncovers recurrent patterns in gene families

As some genes with thesaurus mutations already have established links to cancer, we began an exploration of hits by considering the overlap of all thesaurus genes with the cancer gene census^26^. Our pipeline detected thesaurus mutations in the coding sequences of 35 census genes (Figure 4a) and the untranslated or promoter regions of 29 more (Figure S11). In four of these genes *(NUTM2A, NUTM2B, SSX2, SSX4),* thesaurus mutations comprised their entire mutational load. This is consistent with these genes being recorded in the census because of translocations and fusions, which are detected by algorithms other than somatic substitution calling. Thesaurus substitutions in these cases provide a complementary set of mutation events. For other genes, thesaurus mutations raised the cohort frequency from a non-zero base. The proportion of samples affected exclusively by thesaurus mutations ranged from a few percent (e.g. 5% for *PIK3CA,* 2% for *ROBO2)* up to a large majority (e.g. 88% for *RGPD3).*

**Figure 4.**
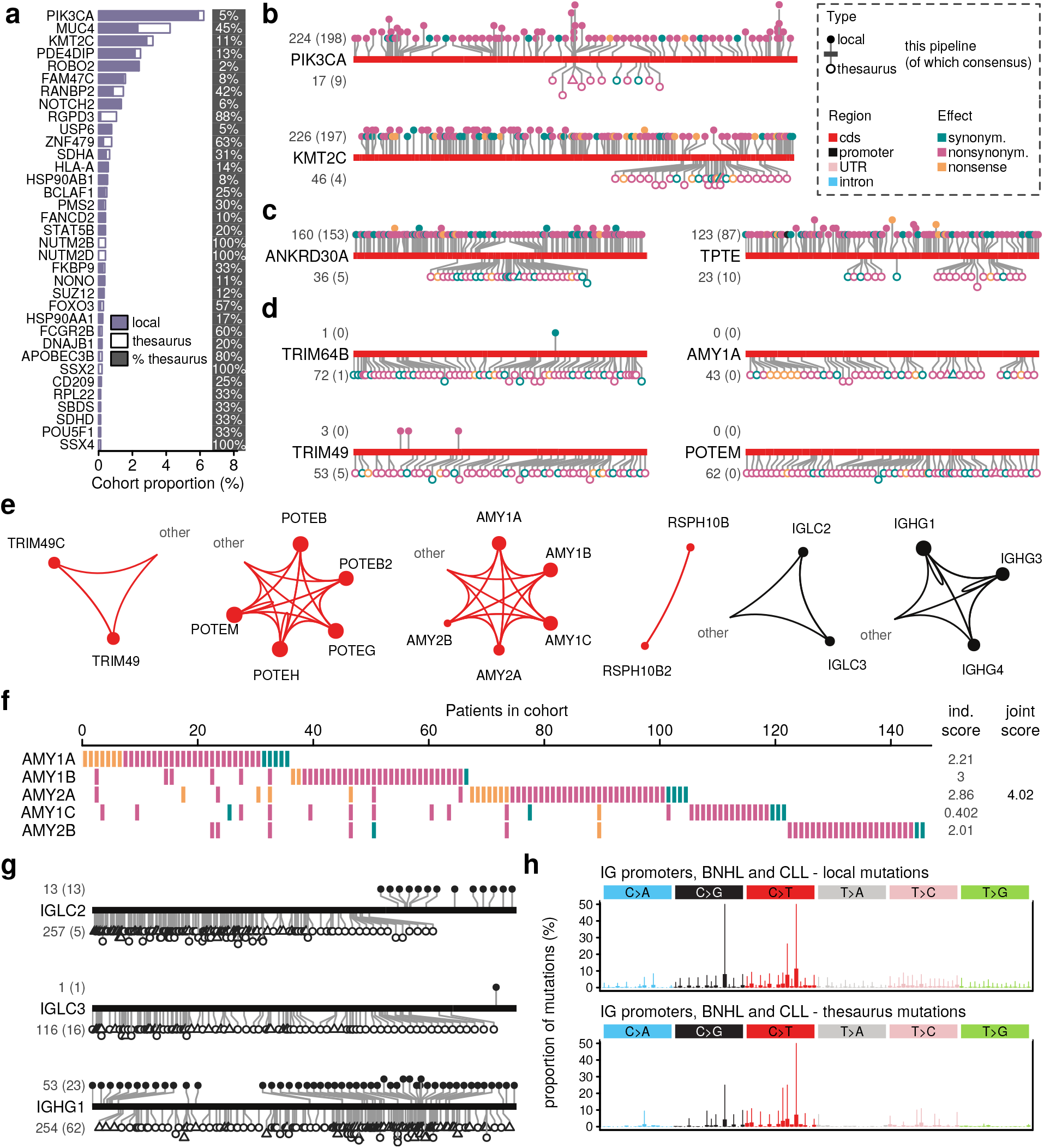
Thesaurus mutations in gene families. **(a)** Thesaurus mutations in coding sequences of the cancer gene census. Percentages on right indicate the proportion of patients that carry exclusively thesaurus mutations. **(b)** Distribution of mutations along the cds of two known cancer genes. **(c)** Analogous to (b), genes carrying both local and thesaurus mutations, but not part of cancer census. **(d)** Analogous to (b) with genes carrying mostly, or exclusively, thesaurus mutations. **(e)** Dominant linking patterns between selected genes. Genes and links are colored according to region type, e.g. cds and promoters. Links from one region type to another, e.g. from cds to UTR, are grouped into a label ‘other’, **(f)** Cohort summary of mutations in one gene family. **(g)** Distribution of mutations along sequences upstream (promoters) of selected immunoglobulin (IG) fragments. **(h)** Mutation profiles in promoter sequences of B-cell non-Hodgkins leukemia (BNHL) and chronic lymphocytic leukemia (CLL) consistent with somatic hypermutation by deamination.

To gain more insight into where the mutations lay along the gene structure, we visualized variant-level results along the gene sequences, splitting the results by mutation type and comparing with the PCAWG mutation calls (Figure S12, S13). For *PIK3CA* and *KMT2C,* two of the genes with the highest mutation load, our pipeline detected 13% and 15% additional simple substitution events compared to the PCAWG set (Figure 4b). This is broadly consistent with our previous comparisons with the consensus and can in part be attributed to technical differences in the pipelines^17^. Thesaurus mutations were located in 630bp and 2.63kb segments of their coding regions, respectively. There was a significant enrichment compared to the PCAWG calls (Fisher tests, *PIK3CA,* p=7×10^-4^, *KMT2C,* p=2×10^-12^), and, especially for *KMT2C,* the new mutations filled a noticeable gap in the distribution of simple variants.

Outside of established cancer genes, most genes that harbor thesaurus mutations also contain at least some simple substitutions. Examples with heavy mutation burden include *ANKRD30A,* an ankyrin-repeat containing gene linked with breast cancer^27^, and *TPTE,* a phosphatase linked to the *PTEN* pathway^28^ (Figure 4c). Given their high mutation load and interactions with cancer pathways relevant to patient stratification schemes^29^, thesaurus mutations that fill gaps in their mutation profiles offer direct opportunities to test their translational relevance.

While thesaurus mutations constitute a minority of hits for most genes, they represent the dominant class for others (Figure 4d). Among genes highlighted by our z-score analysis and also by alternative prioritization methods, *TRIM49* and *TRIM64B,* two members of the tripartite motif family of proteins were prominent with mutations along their entire gene body. This family is involved in innate immunity, autophagy and carcinogenesis^30^. *AMY1B* encodes an amylase isoenzyme typically expressed in the salivary gland and the pancreas and is embedded in a region of variable copy-number. It may influence metabolism, but its high mutation burden may also be a corollary of the genomic fragility of its surrounding genomic region^31^. *POTEM* is another gene with an ankyrin domain with heavy mutation burden. The gene family has been discussed in studies of expression-based biomarkers^32^.

Each of the thesaurus mutations in our dataset is annotated by a link to at least one alternative genomic site with a similar surrounding sequence. These links describe ambiguities in assigning the location of the individual somatic events. True to the ubiquity of non-uniqueness across the genome, we found mutations in coding sequences linked to a variety of targets, including untranslated, intronic, promoter, or untranslated regions of other genes. Mutations in PIK3CA, for example, linked to an untranslated region of a pseudogene. However, we observed that links often interconnected genes from the same family (Figure 4e). These cliques prompted us to consider mutation load across several related genes. Using the amylase gene family as an example, the detected mutations across five genes affected 3.1% of non-hypermutated samples in the cohort. Samples affected by the individual genes were to a large extent non-overlapping (Figure 4f), ruling out the possibility that the mutation set is dominated by double-counting. When we modeled the entire gene family as a single entity in our quantile regression model and z-scoring, the score for the family rose from 3.01 for AMY1A alone to 4.04 for the gene group. Analogous patterns, with variability on the degree of effect on the joint scores, occurred in other families (Figure S14-S22).

Among genes highlighted due to mutations in promoter regions were several members of the immunoglobulin (IG) family (Figure 4g, Figure S23, S24). The immunoglobulin locus undergoes hypermutation during B-cell maturation through cytidine deamination^33^ and genomic rearrangements. Consistent with the role of this process in antibody diversification and immunity, mutations associated to IG gene fragments were enriched in leukemias and lymphomas. Among sequences upstream of all IG gene segments, thesaurus mutations represented 19.7% of all variants in those cancers. Furthermore, the trinucleotide mutation profiles were dominated by C>T substitutions and were consistent with reported sequence hotspot patterns^33^ (Figure 4h). This suggests that thesaurus annotation reliably detects these non-germline events and can thus inform translational approaches that use immune signatures.

## Discussion

The structure of the human genome has been shaped by its evolution, including by duplications and rearrangements. This history leaves a substantial portion of the genome to appear nonunique at the scale of short reads used by high-throughput sequencing studies. As long-and linked-read sequencing protocols become established and widespread, these regions will become accessible for direct analysis^19^. However, existing datasets, including efforts to sequence whole genomes of pan-cancer primary tumors^1^ and metastases^34^, already offer opportunities to evaluate somatic events in these non-unique regions.

When specific sequences are of interest *a priori,* targeted approaches can perform re-analysis of subsets of sequencing data. This analytic strategy is effective when there are complex rearrangements as in the case of the HLA locus^35^, or sequences are present in a large number of copies as in the case of transposons^36^. However, the human genome also carries areas that are almost exactly duplicated due to recent evolutionary events and are present in fewer than ten copies. Such regions can be studied in a systematic manner through a technique that links specific genomic positions and provides information about clusters based on multi-locus alignments^12^. Our pipeline collects multi-locus annotations and leverages high-quality mutation calls from PCAWG to train a machine learning model. The resulting calls for somatic substitutions provide a first systematic summary of the mutation events in non-unique regions, at a genome-wide scale, across several cancer types.

The landscape of thesaurus-annotated mutations covers over 1700 coding genes, 4500 promoters, and thousands of other functional elements. The mutation burden in these regions is consistent with known properties of cancer types and trinucleotide substitution patterns that characterize underlying molecular mechanisms are concordant as well. Mutation rates are in line with those in genes with unique sequence, and outliers are affected to an extent comparable to established cancer genes. These hits provide tantalizing leads toward a more complete picture of mutational processes in cancers.

Our analyses of cohort mutation rates, regional recurrence and hotspots, cancer-type specificity, and co-occurrence are a first-pass summary of the patterns in these data. Indeed, mutational processes are modulated, directly or indirectly, by a myriad of factors that include nucleotide content, chromatin accessibility, gene expression^22^. While methods developed in these areas provide guidance for more refined analyses, they rely on auxiliary data as model covariates. In the context of non-unique regions, these covariates, as they are often acquired through short-read sequencing, are likely to suffer biases related to sequence uniqueness^37^. A careful examination of those covariates in the non-unique genome is a critical step toward better understanding of the statistical and functional importance of the uncovered mutational landscape.

Beyond somatic substitution events, cancer genomes also suffer other types of mutations, including small insertions and deletions and regional copy-number changes^4^. Such events are fewer in number than SNVs but can have more profound functional consequences. Our dataset carries evidence that such events occur in genes with non-unique sequences. Our analysis, however, does not include them because of current limitations in thesaurus annotation. It is thus clear that, despite uncovering thousands of additional somatic events, more still remains hidden. Similar considerations are also relevant for comparative genomics of germline variants and their relation to rare diseases^38^. The challenges in detecting these events are technical, but can be overcome with careful strategies for variant comparison^39^, and would help refine views on the genomics of disease.

## Material and Methods

### Cancer cohort

Data for cancer samples and matching normal tissue were obtained through the Pan-Cancer Analysis of Whole Genomes (PCAWG) consortium^1^. The dataset consisted of 2,658 samples from 38 distinct cancer types (22 organ systems) and 47 consortium projects. All alignments were used in their original formats as provided by the consortium. Briefly, all samples consisted of reads aligned to the hs37d5 reference build of the human genome using bwa-mem^40^.

A random set of 237 normal-tissue samples reported not to be contaminated by tumor cells were selected to form a panel of normals. These samples originated from all 38 cancer types. The panel thus captures heterogeneity of human populations, although it cannot be treated as a true representative of all genetic variation.

### Annotations

All calculations were performed against the hs37d5 genome reference build, including decoy chromosomes. Gene annotations were obtained from GENCODE, release 19^41^. Classifications of known repeat elements were downloaded from the UCSC genome browser hub.

### Variant calling, thesaurus annotation, and candidate prioritization

An initial set of variants were called from bwa-aligned sequencing data using Bamformatics (v0.2.5) (https://github.com/tkonopka/Bamformatics). Settings were left at their defaults, except for argument --minmapqual 0, which instructs the software to use all primary read alignments irrespective of mapping quality.

Variants detected in samples included in the panel of normals were aggregated into a single table. The frequency of each variant in this panel was kept as a proxy for population frequency.

Variants from tumor samples were annotated using GeneticThesaurus (v0.2.1) (https://github.com/tkonopka/GeneticThesaurus). Customized settings included --minmapqual 0, which instructs the software to use all primary aligned reads irrespective of mapping quality, and settings --many 20 --toomany 100, which limit thesaurus links to a smaller number than set by default. The thesaurus annotation process was provided access to alignment data for matched normal samples. Output from this stage included tables linking variants to related sites in the genome, as well as tables associating each variant in the tumor with features such as allelic frequency, coverage, and analogous features from the matched normal. These tables included data based only on the called variant position as well as data pooling information from all thesaurus-linked sites.

Following the variant annotation by the GeneticThesaurus software, variants from each tumor were compared to the panel of normals. Items present at a frequency greater than 1% in normal samples were deemed to consist of common germline variants or sequencing artefacts and were excluded from downstream analysis. Prioritized candidates were further annotated with features from the CIGAR strings of aligned reads using custom scripts.

### Machine learning to detect somatic mutations

A set of 300 tumor samples and their matching normal controls were selected at random for machine-learning. The set contained representatives from all the major PCAWG histologies. The set was split into training, testing, and validation sets with 150, 50, and 100 samples, respectively. Separately, PCAWG mutation calls for the same samples were assembled into a truth set. Importantly, the PCAWG mutation calls were filtered using the same panel-of-normals frequency filter used on the candidate sites and then trimmed further to remove items that were not present in the variant candidates. These steps ensured that the candidates and the truth set are consistent, and that all items in the truth set could in principle be obtained from non-missing features in the candidate data. Both the candidate and the truth dataset were restricted to sites with a PASS filter code in the candidate data, i.e. to those sites not linked to any additional locations via thesaurus links. This ensured that the identification of somatic mutations among the candidates could be determined by technical features of the sequencing data and biological aspects of the tumor and was not confounded by aspects related to mappability.

Machine learning models were trained using xgboost^42^ – an algorithm based on random forests – with default settings except when stated. To explore the effect of data quantity on classification performance, a series of models were trained based on an increasing number of tumor samples. The samples used in each model were selected with a stochastic procedure that attempted to use distinct samples in replicates. Once the tumor samples were selected, models were trained using all the data from those tumors. After training, all models were evaluated against the entire test set using false-discovery (ratio of false calls among all positive calls) and false-negative rates (ratio of missed calls among all calls in the truth set). After obtaining a satisfactory set of hyperparameters, a final model was trained using the entire set of 150 training samples and reevaluated on an independent validation set.

Feature importance in the final classifier was assessed using a bootstrapped dropout-loss procedure. This procedure subsamples the testing dataset, permutes values within individual features or certain groups of features, and assesses how the predictions on the adulterated data compare with predictions based on the original data. The downsampling and permutation procedure was repeated 100 times and average dropout-loss values reported.

This model was then applied to call somatic single nucleotide variants on the entire set of candidate sites irrespective of filter code, i.e. on unique and non-uniquely mappable sites. Importantly, for features that can be affected by mappability, the values provided to the classifier were those estimated by the GeneticThesaurus annotation procedure, i.e. averages over all sites linked by the thesaurus. By construction, this procedure ensured that in well-mappable areas, the classifier functioned in the same way as during training. In non-unique regions, the classifier made predictions from data informed by multiple locations in the genome.

### Modelling mutation load

Called mutations were classified into two groups – local and thesaurus – based on whether a site was associated with an alternative location via a thesaurus link. The mutation load in each sample was defined as the total number of positions for each type. Cases where mutations were called at more than one site and linked together via a thesaurus annotation were treated as single events; the site with the higher allelic frequency was marked as primary and taken forward for subsequent analysis. The relation between the mutation loads associated with local and thesaurus filter codes was modeled using a power-law equation

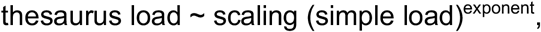

where scaling and exponent are free parameters. Under an assumption that thesaurus mutations appear via the same mechanisms as simple mutations, power would be equal to unity and proportionality would indicate the relative size of non-unique and unique genomic regions. The model can be reformulated as, log(thesaurus load) ~ scaling + exponent log(simple load). The free parameters were solved using simple regression.

### Linked-read sequencing

A sample from a patient with a malignant peripheral nerve sheath tumor was selected for sequencing. The patient provided their written and informed consent to provide samples for this study, which was approved by the National Research Ethics Service (NRES) Committee Yorkshire and The Humber – Leeds East (15/YH/0311).

High molecular weight DNA was extracted from frozen tissue according to the 10x genomics protocol (CG000072). In short, 500 μl of cold nuclei isolation buffer was added (sigma pure prep lysis buffer (NUC201-1KT), 1mM DTT, 10% Triton X-100) to a small piece of frozen tissue in a 1. 5 ml tube (and homogenized by moving a pestle up and down 10-20x). After settling, the supernatant was transferred to a new tube and centrifuged for 5 min at 500xg (4°C). The supernatant was removed without disturbing the pellet and the following was added: 70 μl of cold PBS, 10 μl of proteinase K and 70 μl of Digestion buffer (20mM EDTA, 2nM Tris-HCl, 10mM N-laurylsarcosine sodium salt and water). The pellet was dislodged by tapping the tubes lightly and then left to rotate for 2h at 20°C. Tween-20 was added to a final concentration of 0.1% and pipette mixed 5 times. An equal amount of 1x SPRIselect reagent (Beckman Coulter B23317) was added followed by rotation for 20 min. The beads were then washed twice with 70% ethanol and resuspended in 50 μl sample elution buffer (Qiagen AE buffer with 0.1% Tween 20). After incubation at 20°C for 5 min, the beads were put on a shaker at 1400rpm for 3 min (25°C) to elute the DNA. The samples were quantified using the Qubit. Linked reads libraries were generated using the 10x Chromium, following the manufacturer’s instructions. The library was sequenced using paired-end 150bp reads with an 8bp index on a single lane of an Illumina HiSeq X.

### Validation through linked-read sequencing

BCL files were processed and demultiplexed to FASTQ files using bcl2fastq v2.20.0. Reads were mapped to the hs37d5 reference build, de-duplicated and filtered using the LongRanger (v2.2.2) WGS pipeline with the --somatic flag. This pipeline leverages the Chromium molecular barcodes and GATK v4.0.8.1 to call and phase single nucleotide variants, indels, and structural variants. Overall, a total of 1,538,338 GEMs were generated, containing on average 589kb of DNA with an average size of 86kb, each producing a median of 47 linked reads for a final 38x depth of coverage and 32x median depth at mutated sites.

The reported variants were compared to the calls from the machine learning approach on a standard short-read WGS dataset of the same tumor sample. Simple variants called from the short reads were declared validated if they were part of the PASS mutations in the linked reads. Thesaurus calls were declared validated if they were part of PASS mutations in the linked reads, or if their thesaurus-linked sites were part of the linked-read dataset. This approach allows for ambiguity in placing the mutation location based on short-read data, but does not inflate detection rates^12^.

### Mutation trinucleotide profiles

Trinucleotide contexts were extracted for all called mutations in the cohort. These neighborhoods were used to assign each substitution mutation to one of 96 categories as previously described^14^. Counting the number of mutations of each type produced two profiles – one based on simple mutations in unique genomic regions and one based on mutations with thesaurus links.

For correlation analysis, the simple and thesaurus profiles were treated as 96-dimensional vectors and correlation was evaluated using the Spearman method. Because mutation profiles are degenerate when the overall number of mutations is small and constrained to non-negative counts, statistical significance was estimated by simulation. For a given profile of thesaurus mutations, 10,000 random profiles were generated with an equivalent number of mutations. The Spearman correlation value between the local and thesaurus profiles were compared to the distribution of correlations between the local and simulated profiles. The procedure provided approximations to p-values that were sufficiently precise to determine significance at nominal and multiple-testing adjusted levels.

For visualization of similarities of the mutation profiles, count-based mutation profiles were adjusted using allele-frequency data. This adjustment provided weighting toward well-measured mutation instances and avoided degenerate comparisons based on integer counts. Mutation profiles based on simple mutations were sum-normalized and embedded into a two-dimensional space using UMAP^43,44^, a dimensional reduction technique, using a euclidean distance metric. Following generation of the embedding based on simple mutations, the resultant model was used to predict the position of allele-frequency-adjusted profiles based on thesaurus mutations.

### Modeling of mutation frequency

Mutation frequencies were computed by counting the number of samples (patients) with at least one somatic mutation in regions of interest. In order to avoid counts being inflated by likely passenger mutations, hyper-mutator samples were identified and removed from this calculation and subsequent modeling. Hyper-mutator status was set if a sample contained more than 300 mutations in coding regions, a procedure previously described in other studies of driver mutations^22,37^. In practice, we omitted 198 samples (7.5% of the cohort) from the mutation frequency analysis.

Mutation frequencies were assessed on non-overlapping genomic regions labeled as coding (cds), intronic (intron), untranslated (UTR), promoter, or intergenic. Each region was associated with a genomic length, a frequency based on simple mutations alone, a frequency based on thesaurus mutations alone, and a frequency based on both simple and thesaurus mutations.

Modeling of the relation between region size and mutation frequency was performed using quantile regression. The model used logarithmically transformed frequency and region size,

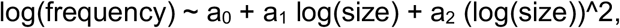

With a_0_, a_1_, and a_2_ as free parameters. The model quadratic term allows some nonlinearity in the relationship between size and frequency, which is required to allow the growth in frequency to taper for very large regions. Quantile regression with this model was performed at the 50% percentile to describe the primary trend, and at 5%, 25%, 75%, 95% levels to obtain intervals of variability. After fitting the parameters, each genomic region was associated with an expected mutation frequency and an interquartile interval.

Quantile regression based on a linear equation is guaranteed to produce fitted models that preserve ordering of percentiles, e.g. with a model at quantile of 75% always yielding larger values than at 50%. This property is not guaranteed for models with higher-order terms. Predictions from the fitted models were thus adjusted post-hoc. Furthermore, the interquartile interval was forced to correspond to at least 1/N, with N being the number of samples in the modeled cohort. All model predictions were restricted to the unit interval, [0, 1].

### Identifying driver genes with dndscv

The dN/dS ratio was computed through maximum likelihood estimates across trinucleotide contexts using dndscv^22^. After removing hypermutator samples with more than 300 coding mutations, we ran dndscv on the pan-cancer cohort for *de-novo* discovery of candidate cancer genes. We used the pooled set of thesaurus and simple mutations, but we did not correct for epigenetic covariates, as these covariates are not annotated for non-uniquely mapping regions.

### Cancer type specificity

Specificity of mutations was quantified using information entropy, defined via Shannon’s formula,

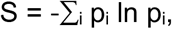

with the sum running over cancer types. Probabilities p_i_ for each cancer type were defined as p_i_ = (c_i_ + *ε*)/ Σ_j_(c_j_ + *ε*) with c_i_ being the count of patients (samples) carrying a mutation in the region and the constant # being a pseudocount regularization. For the regularization, we used a value of 1 for coding sequences and adjusted values for promoters, UTRs, introns, and intergenic regions proportionally to the median region length. Because entropy is high for quasi-random configurations and low for configurations peaked on one bin, visualizations of specificity were performed using changes in entropy, ΔS. These were defined by subtracting S from the entropy of a hypothetical region with null counts,

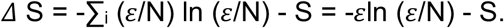

The quantity gives mutation patterns with a high number of patients from a single cancer type higher values than cancer-type-agnostic patterns. It also assigns values near zero to all infrequently-mutated regions.

## Acknowledgements

This work is supported by The Francis Crick Institute, which receives its core funding from Cancer Research UK (FC001202), the UK Medical Research Council (FC001202) and the Wellcome Trust (FC001202). M.T. is supported as a postdoctoral fellow by the European Union’s Horizon 2020 research and innovation program (Marie Skłodowska-Curie Grant agreement no. 747852-SIOMICS). J.D. is a postdoctoral fellow of the Research Foundation – Flanders (FWO). AMF is a NIHR senior investigator, and is supported by the National Institute for Health Research, UCLH Biomedical Research Centre and the CRUK Experimental Cancer Centre. P.V.L. is a Winton Group Leader in recognition of the Winton Charitable Foundation’s support toward the establishment of The Francis Crick Institute. T.K. would like to thank Damian Smedley. This project was enabled through the Crick Scientific Computing STP and through access to the MRC eMedLab Medical Bioinformatics infrastructure, supported by the UK Medical Research Council (grant no. MR/L016311/1). The Bone Cancer Research Trust funded sample biobanking.

(External File)

**Supplementary Table 1. Summary of z-scores and histology specificity for all genomic regions.**

## References

1. The ICGC/TCGA Pan-Cancer Analysis of Whole Genomes Consortium. Pan-cancer analysis of whole genomes. Nature 578, 82–93 (2020).

2. Alexandrov, L. B. et al. The Repertoire of Mutational Signatures in Human Cancer. Nature 578, 94–101 (2020).

3. Li, Y. et al. Patterns of somatic structural variation in human cancer genomes. Nature 578, 112–121 (2020).

4. Rheinbay, E. et al. Analyses of non-coding somatic drivers in 2,658 cancer whole genomes. Nature 578, 102–111 (2020).

5. Gerstung, M. et al. The evolutionary history of 2,658 cancers. Nature 578, 122–128 (2020).

6. Karimzadeh, M., Ernst, C., Kundaje, A. & Hoffman, M. M. Umap and Bismap: quantifying genome and methylome mappability. Nucleic Acids Res. 46, e120 (2018).

7. Lee, H. & Schatz, M. C. Genomic dark matter: the reliability of short read mapping illustrated by the genome mappability score. Bioinformatics 28, 2097–2105 (2012).

8. Mandelker, D. et al. Navigating highly homologous genes in a molecular diagnostic setting: a resource for clinical next-generation sequencing. Genetics in Medicine vol. 18 1282–1289 (2016).

9. Suzuki, I. K. et al. Human-Specific NOTCH2NL Genes Expand Cortical Neurogenesis through Delta/Notch Regulation. Cell 173, 1370–1384.e16 (2018).

10. Suzuki, H. et al. Recurrent noncoding U1 snRNA mutations drive cryptic splicing in SHH medulloblastoma. Nature 574, 707–711 (2019).

11. Shuai, S. et al. The U1 spliceosomal RNA is recurrently mutated in multiple cancers. Nature (2019) doi:10.1038/s41586-019-1651-z.

12. Kerzendorfer, C., Konopka, T. & Nijman, S. M. B. A thesaurus of genetic variation for interrogation of repetitive genomic regions. Nucleic Acids Res. 43, e68 (2015).

13. Konopka, T. & Nijman, S. M. B. Comparison of genetic variants in matched samples using thesaurus annotation. Bioinformatics 32, 657–663 (2016).

14. Alexandrov, L. B. et al. Signatures of mutational processes in human cancer. Nature 500, 415–421 (2013).

15. Ainscough, B. J. et al. A deep learning approach to automate refinement of somatic variant calling from cancer sequencing data. Nat. Genet. 50, 1735–1743 (2018).

16. Anzar, I., Sverchkova, A., Stratford, R. & Clancy, T. NeoMutate: an ensemble machine learning framework for the prediction of somatic mutations in cancer. BMC Med. Genomics 12, 63 (2019).

17. Garcia-Prieto, C., Valencia, A. & Porta-Pardo, E. The consequences of variant calling decisions in secondary analyses of cancer sequencing data. doi:10.1101/2020.01.29.924860.

18. Ellrott, K. et al. Scalable Open Science Approach for Mutation Calling of Tumor Exomes Using Multiple Genomic Pipelines. Cell Syst 6, 271–281.e7 (2018).

19. Bishara, A. et al. Read clouds uncover variation in complex regions of the human genome. Genome Res. 25, 1570–1580 (2015).

20. Zheng, G. X. Y. et al. Haplotyping germline and cancer genomes with high-throughput linked-read sequencing. Nat. Biotechnol. 34, 303–311 (2016).

21. Lawrence, M. S. et al. Mutational heterogeneity in cancer and the search for new cancer-associated genes. Nature 499, 214–218 (2013).

22. Martincorena, I. et al. Universal Patterns of Selection in Cancer and Somatic Tissues. Cell 173, 1823 (2018).

23. Chen, H. et al. Comprehensive assessment of computational algorithms in predicting cancer driver mutations. Genome Biol. 21, 43 (2020).

24. Araya, C. L. et al. Identification of significantly mutated regions across cancer types highlights a rich landscape of functional molecular alterations. Nat. Genet. 48, 117–125 (2015).

25. Bailey, M. H. et al. Comprehensive Characterization of Cancer Driver Genes and Mutations. Cell 174, 1034–1035 (2018).

26. Tate, J. G. et al. COSMIC: the Catalogue Of Somatic Mutations In Cancer. Nucleic Acids Res. 47, D941–D947 (2019).

27. Jäger, D. et al. Identification of a tissue-specific putative transcription factor in breast tissue by serological screening of a breast cancer library. Cancer Res. 61, 2055–2061 (2001).

28. Tapparel, C. et al. The TPTE gene family: cellular expression, subcellular localization and alternative splicing. Gene 323, 189–199 (2003).

29. Jamaspishvili, T. et al. Clinical implications of PTEN loss in prostate cancer. Nat. Rev. Urol. 15, 222–234 (2018).

30. Hatakeyama, S. TRIM Family Proteins: Roles in Autophagy, Immunity, and Carcinogenesis. Trends in Biochemical Sciences vol. 42 297–311 (2017).

31. Usher, C. L. et al. Structural forms of the human amylase locus and their relationships to SNPs, haplotypes and obesity. Nat. Genet. 47, 921–925 (2015).

32. Barger, C. J. et al. Expression of the POTE gene family in human ovarian cancer. Sci. Rep. 8, 17136 (2018).

33. Teng, G. & Papavasiliou, F. N. Immunoglobulin somatic hypermutation. Annu. Rev. Genet. 41, 107–120 (2007).

34. Priestley, P. et al. Pan-cancer whole-genome analyses of metastatic solid tumours. Nature vol. 575 210–216 (2019).

35. McGranahan, N. et al. Allele-Specific HLA Loss and Immune Escape in Lung Cancer Evolution. Cell 171, 1259–1271.e11 (2017).

36. Rodriguez-Martin, B. et al. Pan-cancer analysis of whole genomes identifies driver rearrangements promoted by LINE-1 retrotransposition. Nat. Genet. 52, 306–319 (2020).

37. Kundaje, A. et al. Integrative analysis of 111 reference human epigenomes. Nature 518, 317–330 (2015).

38. Eichler, E. E. Genetic Variation, Comparative Genomics, and the Diagnosis of Disease. N. Engl. J. Med. 381, 64–74 (2019).

39. Krusche, P. et al. Best practices for benchmarking germline small-variant calls in human genomes. Nat. Biotechnol. 37, 555–560 (2019).

40. Li, H. & Durbin, R. Fast and accurate long-read alignment with Burrows-Wheeler transform. Bioinformatics 26, 589–595 (2010).

41. Frankish, A. et al. GENCODE reference annotation for the human and mouse genomes. Nucleic Acids Res. 47, D766–D773 (2019).

42. Friedman, J., Hastie, T. & Tibshirani, R. Additive logistic regression: a statistical view of boosting (With discussion and a rejoinder by the authors). The Annals of Statistics vol. 28 337–407 (2000).

43. McInnes, L., Healy, J., Saul, N. & Großberger, L. UMAP: Uniform Manifold Approximation and Projection. Journal of Open Source Software vol. 3 861 (2018).

44. Becht, E. et al. Dimensionality reduction for visualizing single-cell data using UMAP. Nat. Biotechnol. (2018) doi:10.1038/nbt.4314.

