## Supplementary Figures for "A pan-cancer landscape of somatic substitutions in non-unique regions of the human genome"

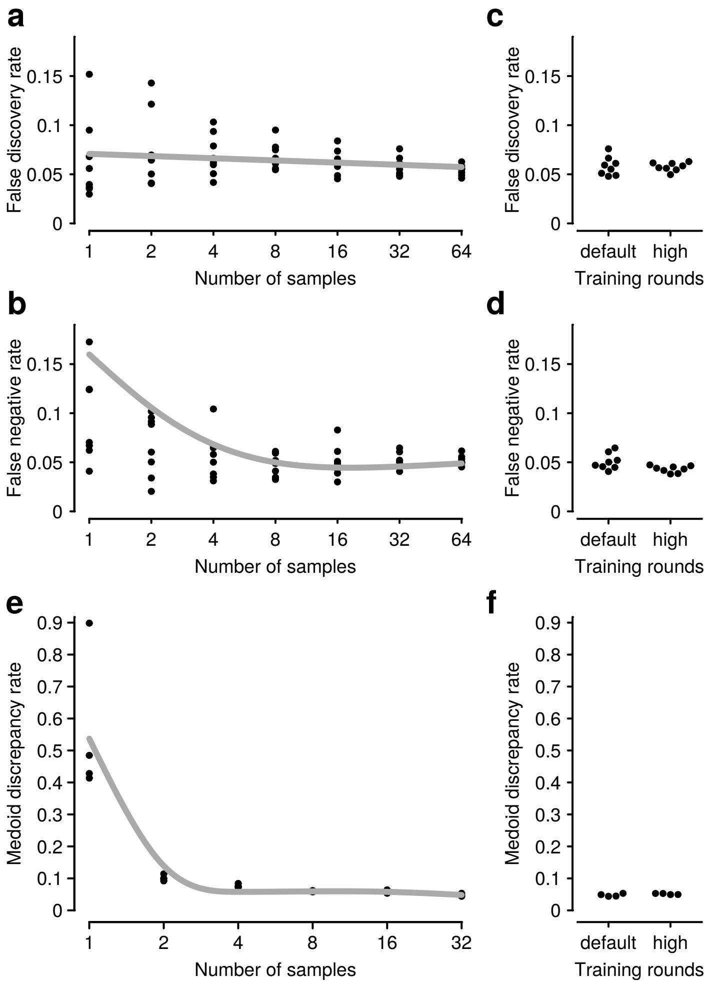


**Figure S1. Training machine learning models to call somatic mutations. (a-d)** A series of models were trained with increasing numbers of samples to explore how volume of training data affects model performance. At each size, 8 models were trained with randomly selected training samples and using 5 xgboost training rounds. All models were evaluated against a fixed test set of 50 samples. At size 32, an additional set of models were trained with 25 xgboost rounds. In all panels, dots correspond to measurements; lines are smoothing curves to summarize trend. **(a)** False positive rate measures new calls with respect consensus calls. **(b)** False negative rate measures propotion of calls in consensus not evaluated by the caller. **(c,d)** Comparison of models of size 32 trained with default and a high number of xgboost rounds. **(e,f)** Another series of models were trained using varying numbers of samples, in batches of size 4, so that each batch used non-overlapping sets of training data. **(e)** Medoid discrepancy rate measures the average difference between calls and the batch medoid; it thus captures consistency between models trained on different samples rather than ability to reproduce a consensus. **(f)** Comparison of models trained using 32 samples with the default and a high number of training rounds.

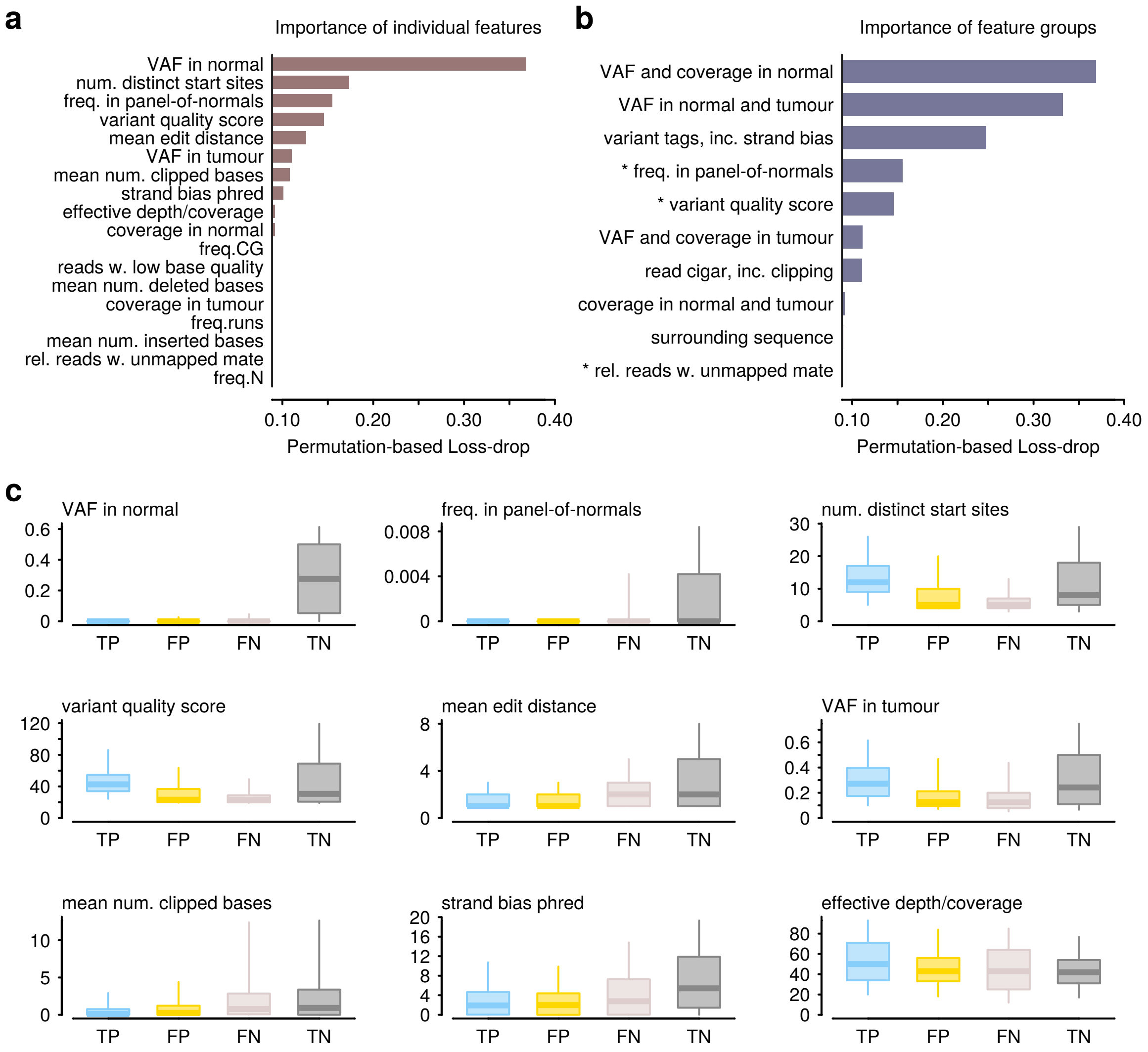


**Figure S2. Feature importance for calling somatic mutations.** A single classifier was trained using 150 training samples with a high number of xgboost rounds, and then evaluated using 50 held-out samples. Feature performance was evaluated by measuring the change in model performance on test data when certain variables were scrambled. **(a)** Dropout-loss after scrambling individual features indicates reveals the importance of individual variables. The vertical line indicates the baseline value of the loss function when all features are included in the model. **(b)** Analogous to (a) but showing the efect of scrambing several variables at once. Variable groups capture combination of related features, and some groups may overlap. Items marked with a (*) represent groups with a single feature and reproduce results from (a). **(c)** Distributions of feature values at positions in the test set classified as true positives (TP), false positives (FP), false negatives (FN), and true negatives (TN) with respect to PCAWG consensus calls. Thick lines represent medians; box boundaries convey the interquartile range; whiskers denote 5% and 95% quantiles.

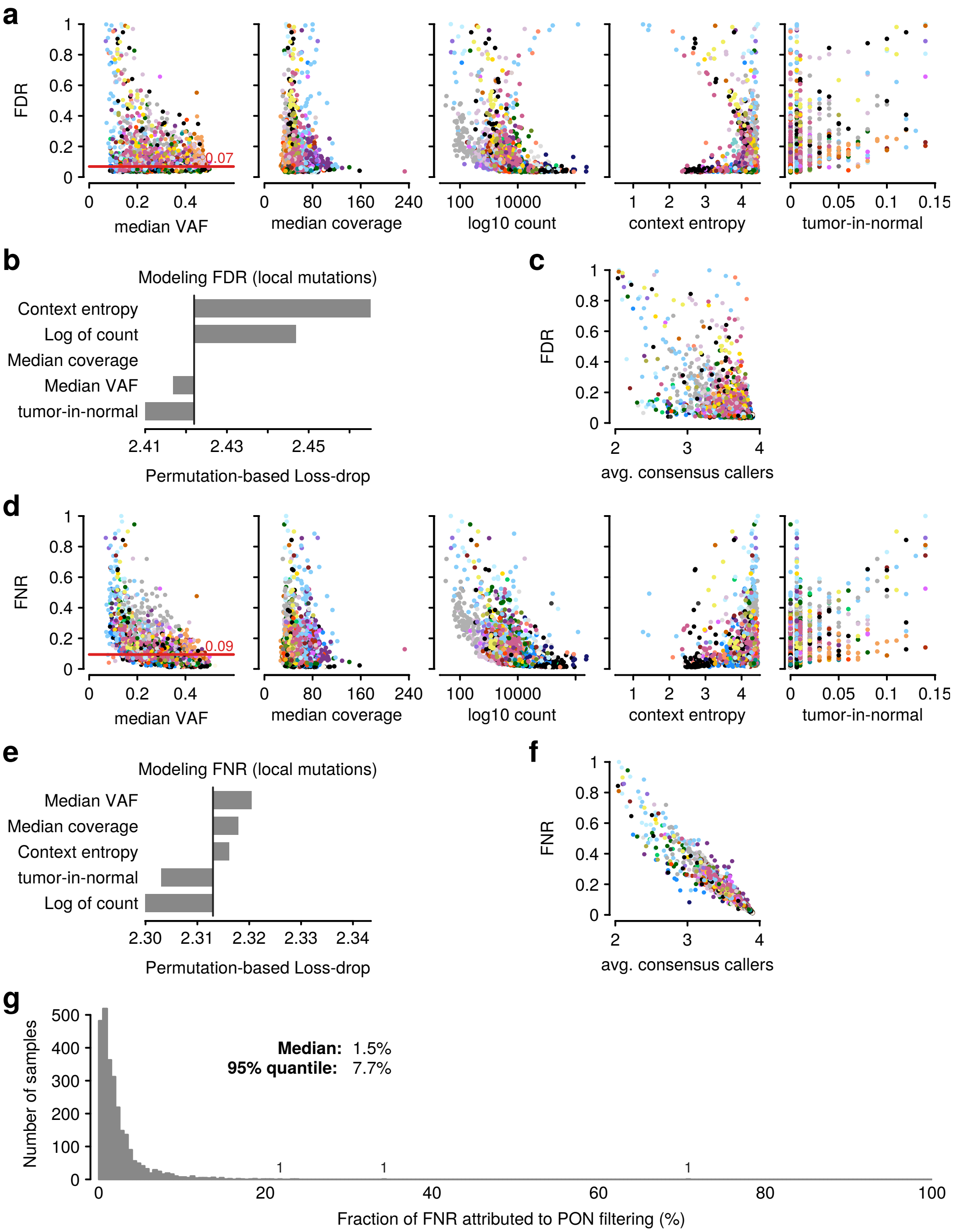


**Figure S3. Factors affecting false discovery rate (FDR) and false negative rate (FNR) compared to the consensus. (a)** Relationships between single features and the FDR. Points represent samples, colored by cancer type. Red line indicates the median FDR in the cohort. **(b)** Feature importance for explaining FDR using a generalized linear model. **(c)** Comparison of FDR against internal concordance in the consensus calls, measured by the averge number of callers that call mutation sites in the sample. **(d,e,f)** Analogous to (a,b,c) for FNR. **(g)** Fraction of FNR that can be attributed to filtering variants because of presence in the panel-of-normals (PON). Numbers along the histograms indicate sample counts in isolated bins.

**a**

| consensus | miss | PASS | thesaurus |
|---|---|---|---|
| yes | FN | TP | thesP |
| no | | FP | thesD |

TPR = TP / (FN + TP + thesP)
FNR = FN / (FN + TP + thesP)
FDR = FP / (TP + FP + thesP)
thesPR = thesP / (FN + TP + thesP)
thesDR = thesD / (FN + TP + thesP + thesD)

**b**

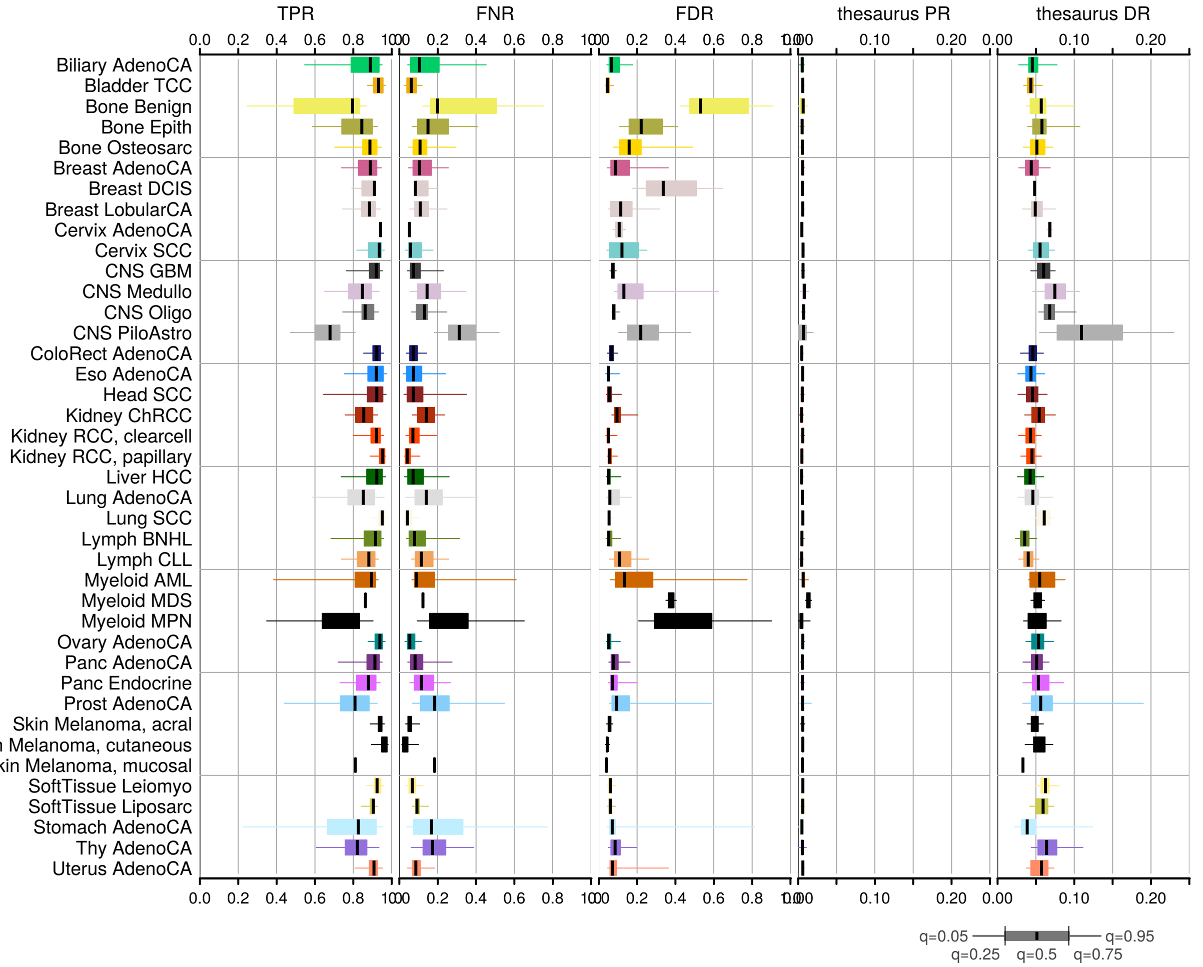


**Figure S4. Comparison of mutation calls with consensus. (a)** Definition of mutation set comparison measures, using the consensus call set as a ‘ground truth.’ FN, false negative; TP, true positive; FP, false positive; thesP, thesaurus positive, i.e. a site labeled as thesaurus linked that is also in the consensus set; thesD, thesaurus discovery, i.e. a novel site labeled as thesaurus linked; TPR, true positive rate; FNR, false negative rate; FDR, false discovery rate; thesPR, thesaurus positive rate; thesDR, thesaurus discovery rate. **(b)** Summary of mutation measures stratified by histology. Each box summarizes a performance measures across samples in the cohort. Box mid-line, edges, and whiskers represent 50%, 25%-75%, and 5%-95% quantiles.

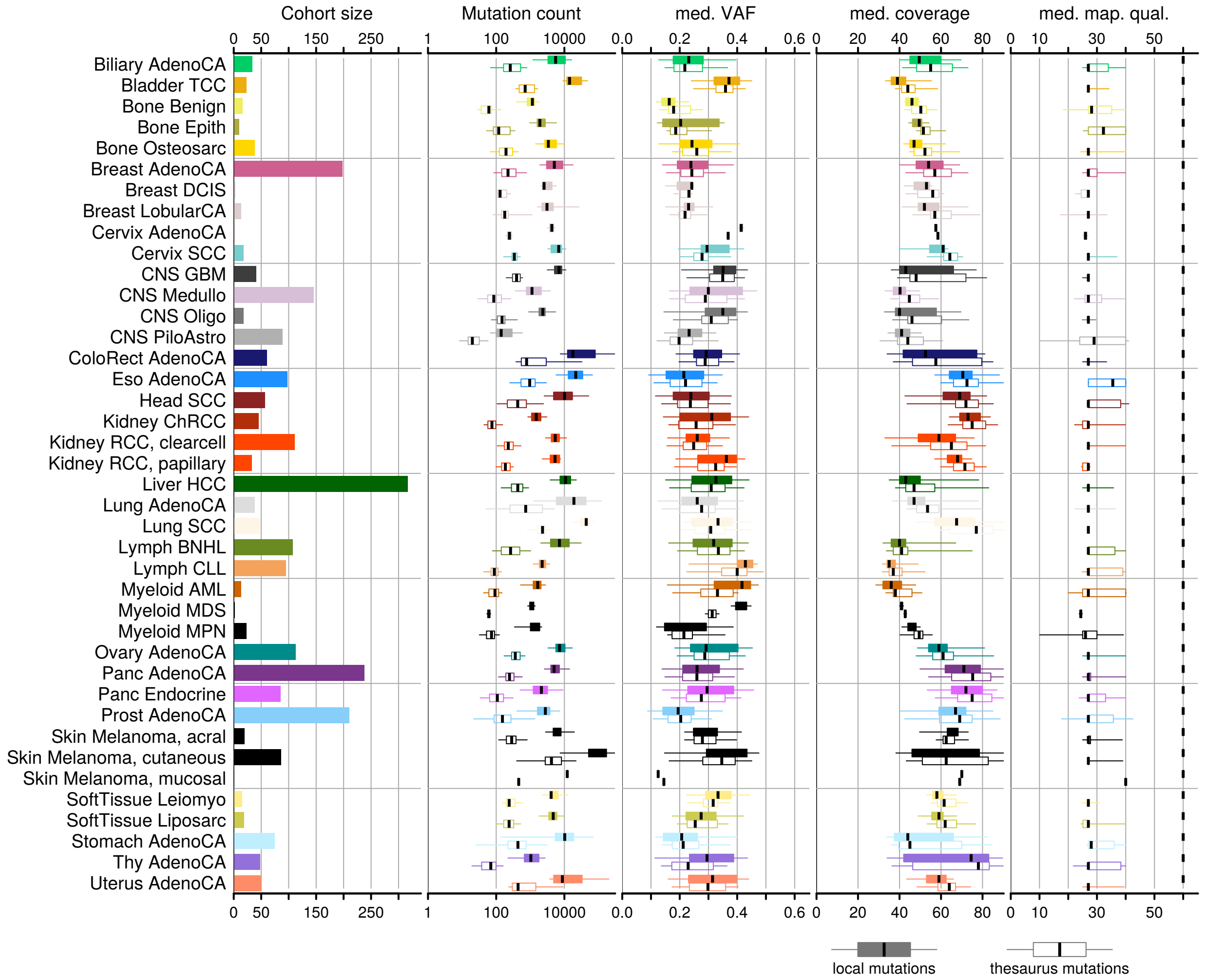


**Figure S5. Characteristics of local and thesaurus mutations.** The left-most panel summarizes the number of samples in each histology cohort. Subsequent panels (left-to-right) summarize distributions of mutation counts, median mapping quality at mutation sites, median variant-allelic frequency (VAF) of the mutant allele, and median coverage at the mutation site. Boxplots display medians (middle line), interquartile ranges (boxes), and 5%-95% quantiles (whiskers). As expected, thesaurus mutations are less numerous than local mutations. Distributions of allelic frequency and coverage are similar for the two groups. Thesaurus mutations are supported by reads with lower mapping quality.

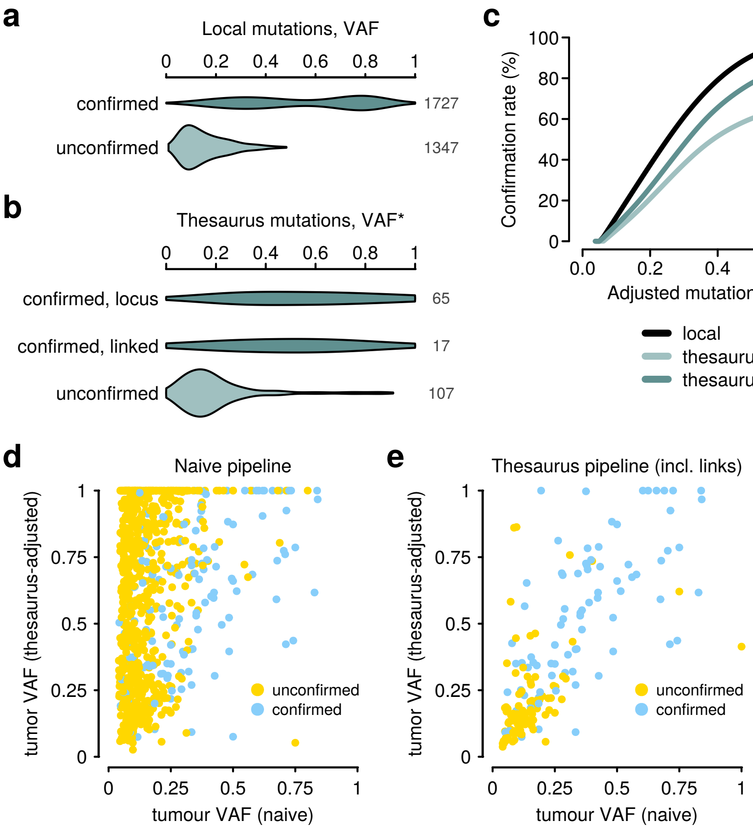


**Figure S6. Comparison of mutations detected in short-read and long-read data.** **(a)** Variant-allele frequency (VAF) of local mutations as measured in short-read data. The mutations are stratified into a group that is confirmed in the long-read data and a group that was not detected in the long-read data. Numbers on the right-hand-side denote the number of mutations in each group. **(b)** Analogous to panel (a), but summarizing thesaurus mutations. One of the stratification groups includes items not detected in the long-read data, but where a linked site was detected in the long-read data. Here, the x-axis is based on thesaurus-adjust allelic frequency. **(c)** Modeling of confirmation rate of mutations as a function of VAF. VAF is measured in the short-read sample using thesaurus adjustment. Lines correspond to spline models with four degrees of freedom. Models consider: local mutations in unique regions, thesaurus mutations with primary flag and evaluated at nominal locus without links (excl. links), thesaurus-filter mutations evaluated using primary flag and links (incl. links). **(d)** Comparison of naive- and theasaurus-adjusted allelic frequency among mutation candidates from a naive pipeline. The validation rate is high among sites with high naive allelic frequency, but the large number of candidates at intermediate frequencies suggest there may be high contamination with germline sites that are 'validated' in the long-read sample. **(d)** Comparison of naive- and thesaurus-adjusted allelic frequncy among candidates from the thesaurus pipeline.

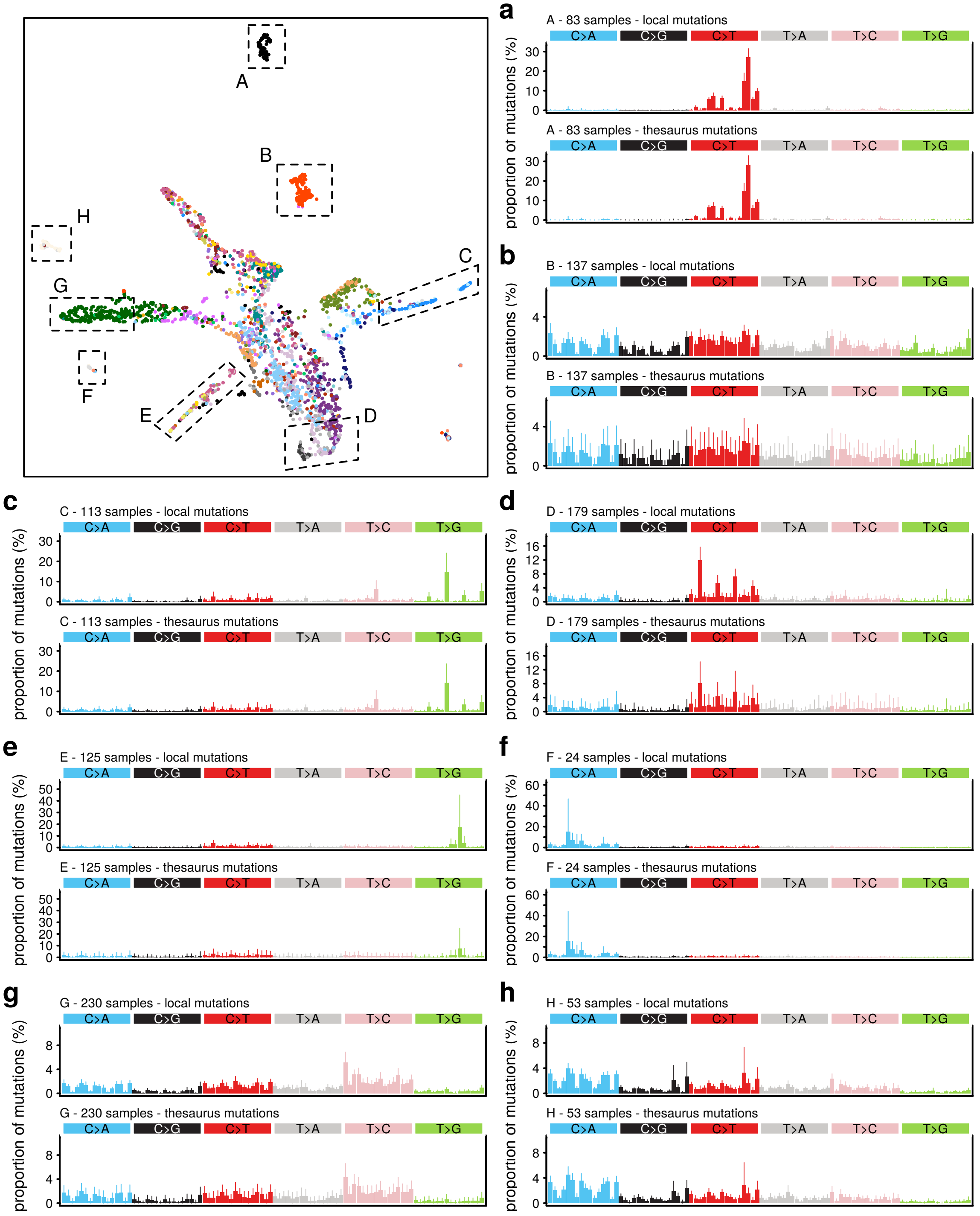


**Figure S7. Mutation profiles in sample groups.** Top-left panel is an embedding of all samples, computed by UMAP using VAF-adjusted mutation profiles of local mutations. Gating highlights manually-selected sample groups. **(a-h)** Panels correspond to gates in the embedding diagram. Top sub-panel displays the average mutation profile based on local mutations (PASS); bottom sub-panel displays average profile based on thesaurus mutations in the same sample group. Error bars denote 5% and 95% intervals.

**a**

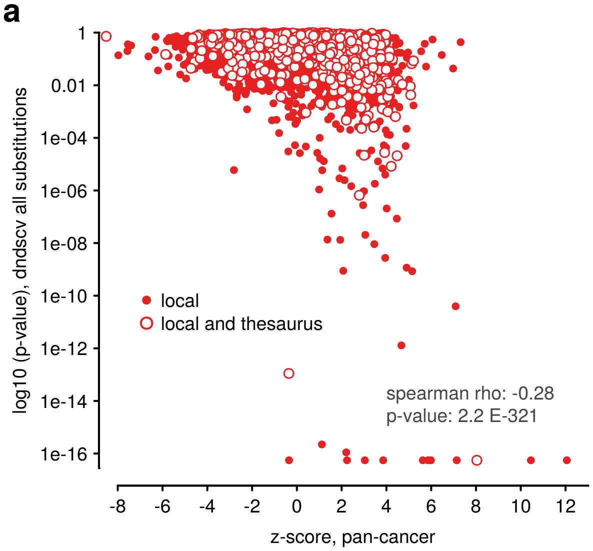


**b**

| gene symbol | thes perc | z score | p value | q value |
|---|---|---|---|---|
| PIK3CA | 5.2 | 8.04 | 5.55e-17 | 8.58e-14 |
| KMT2C | 11.4 | -0.36 | 1.11e-13 | 1.49e-10 |
| HLA-A | 14.3 | 2.79 | 6.55e-07 | 0.000425 |
| ZNF99 | 34.1 | 4.21 | 8.29e-06 | 0.00379 |
| TRIM64B | 100 | 4.48 | 2.08e-05 | 0.00871 |
| AMY1B | 100 | 3 | 2.17e-05 | 0.00874 |
| TRIM49 | 92 | 3.92 | 2.76e-05 | 0.0101 |
| OR4Q3 | 15.4 | 2.89 | 0.000228 | 0.0647 |
| POTEM | 100 | 3.43 | 0.000254 | 0.071 |
| OR11H1 | 100 | 3.48 | 0.000285 | 0.0773 |
| PSG6 | 5.6 | 3.06 | 0.000418 | 0.108 |
| AMY1A | 100 | 2.21 | 0.000471 | 0.115 |
| TRIM49C | 88.9 | 3.08 | 0.000494 | 0.118 |
| POTEH | 97.1 | 4.4 | 0.000642 | 0.143 |
| MUC19 | 0.7 | 5.09 | 0.00427 | 0.47 |
| ANKRD30A | 15.9 | 5.06 | 0.00901 | 0.634 |
| OR4A15 | 3.4 | 4.7 | 0.0108 | 0.661 |
| ZNF208 | 19.3 | 4.53 | 0.0156 | 0.737 |
| FLG | 23 | 5.12 | 0.0658 | 0.93 |
| TPTE | 15.2 | 5.2 | 0.0846 | 0.93 |

**Figure S8. Comparison of gene prioritization schemes. (a)** Comparison of p-values produced by dndscv (all substitutions) against z-scores. Dots represent genes with coding sequence, stratified according to whether they hold only local mutations, or a combination of local and thesaurus mutations. **(b)** Details on a selection of most-significant genes, ranked according to p-value but including genes with high z-scores. The fraction of thesaurus mutations in each gene is indicated as a percentage in column 'thes perc'.

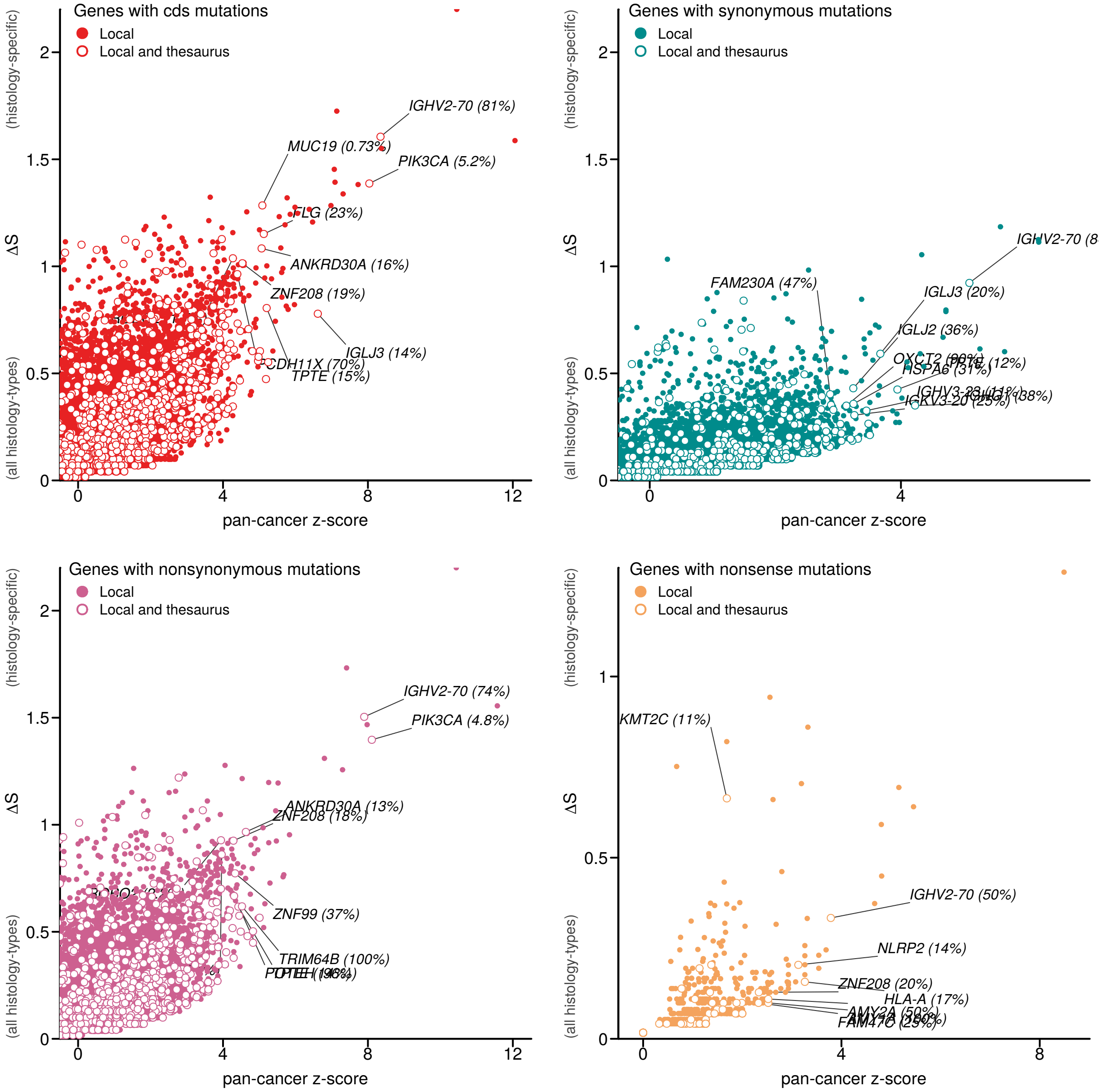


**Figure S9. Specificity of mutations in gene coding sequences.** All panels show the over-representation of mutations in the pan-cancer cohort (z-score, x-axis) and the specificity of the mutation distribution across cancer types (change in entropy, y-axis). One panel visualizes results base on all detected mutations, and subsequent panels use subsets of mutations predicted to have synonymous, non-synonymous, and nonsense effects at the protein level.

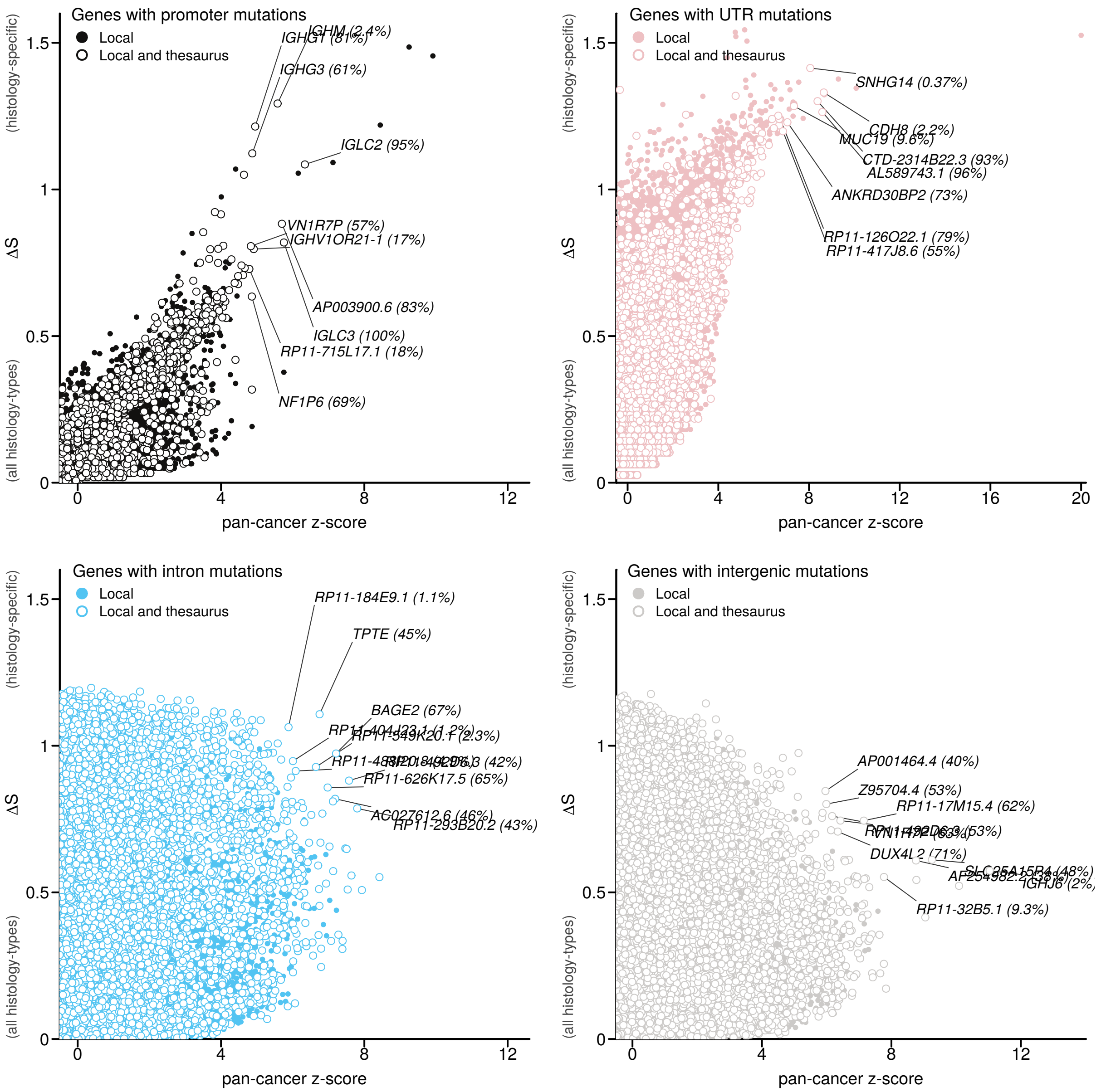


**Figure S10. Specificity of mutations.** All panels show the over-representation of mutations in the pan-cancer cohort (z-score, x-axis) and the specificity of the mutation distribution across cancer types (change in entropy, y-axis). Mutations are split into non-overlapping region sets of promoter regions upstream of coding and non-coding genes, untranslated regions (UTR) associated with coding genes or untranslated region of non-coding genes, intronic regions, and intergenic regions.

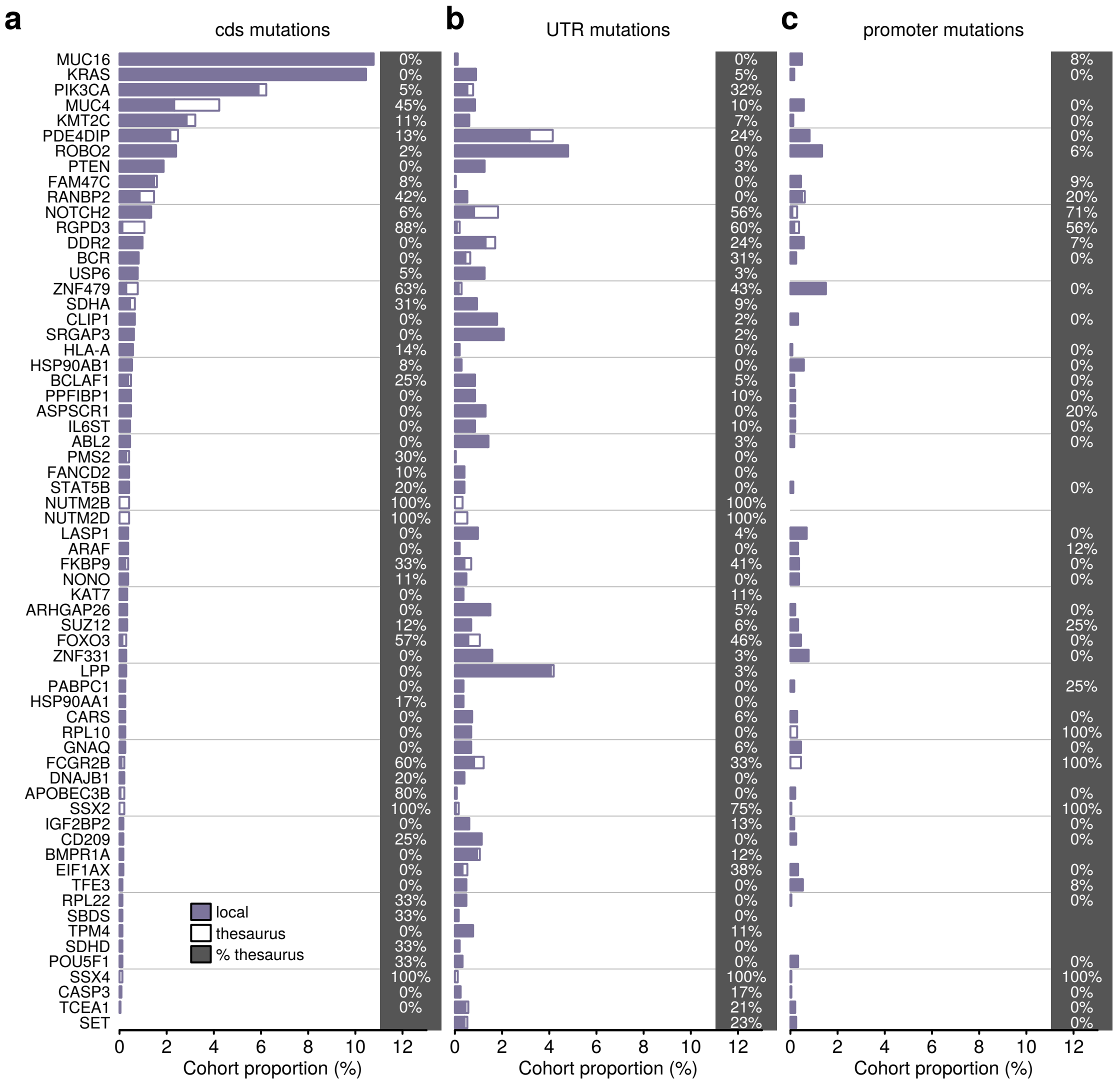


**Figure S11. Genes in cancer gene census that carry thesaurus mutations in cds, UTR, and promoter regions.** Bars are split to convey proportion of cohort that carries a mutation classified as local and thesaurus. Percentages on right-hand-side indicate the proportion of samples that carry exclusively thesaurus mutations. Genes with a proportion of 0% indicates may include thesaurus mutations alongside local mutations.

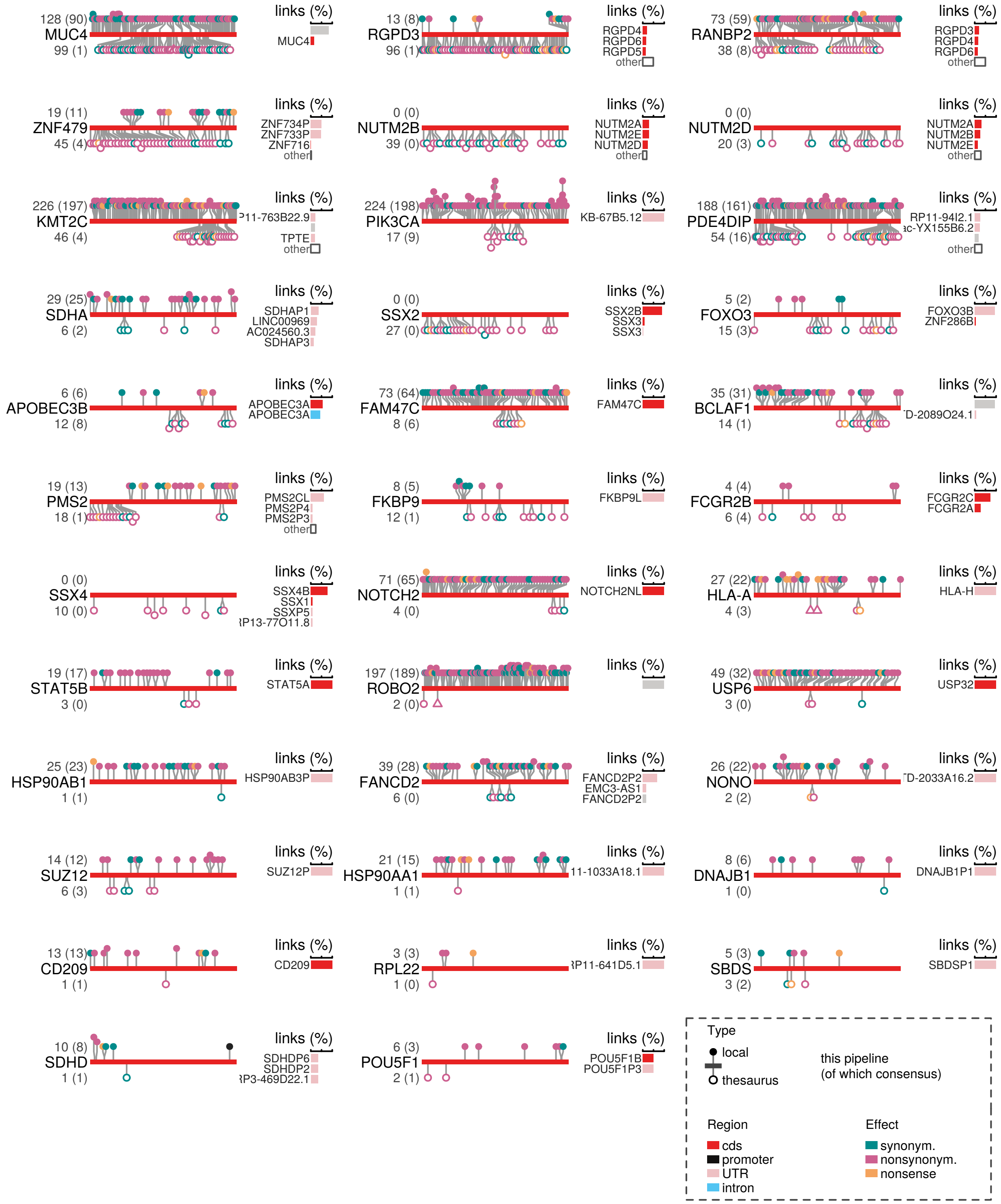


**Figure S12. Mutations along coding regions of genes part of the cancer gene census.** In each example, the coding sequence is represented as a continuous bar; lollipops display the position of mutations along the sequence. Accompanying bar charts indicate genomic locations that thesaurus mutations link toward. Colors of lollipops, regions,

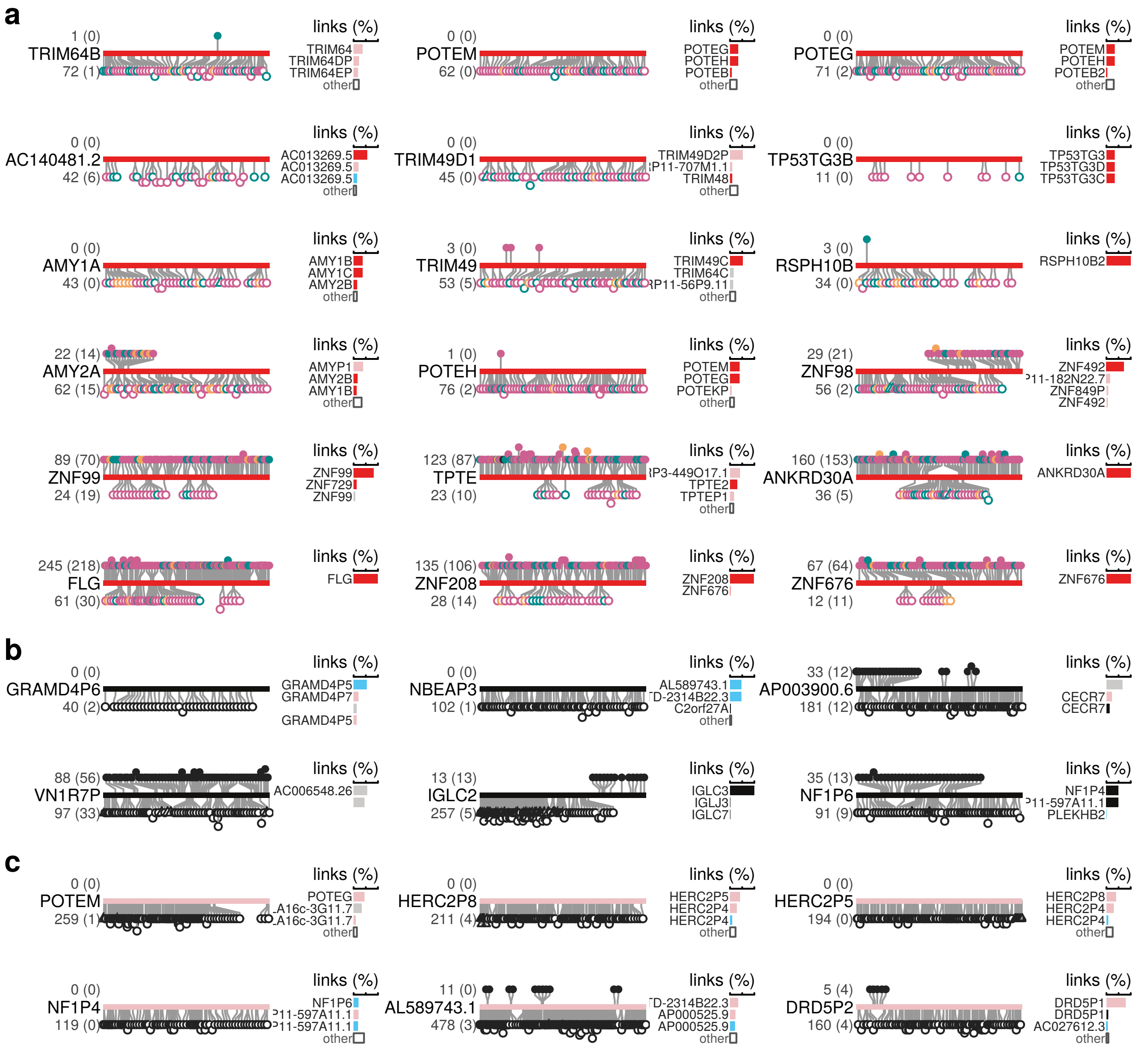


**Figure S13. Selected genes with thesaurus mutations.** Examples are structure as in previous figure. **(a)** Examples of codings regions. **(b)** Examples of promoter regions (sequences upstream of coding sequences). **(c)** Examples of untranslated regions.

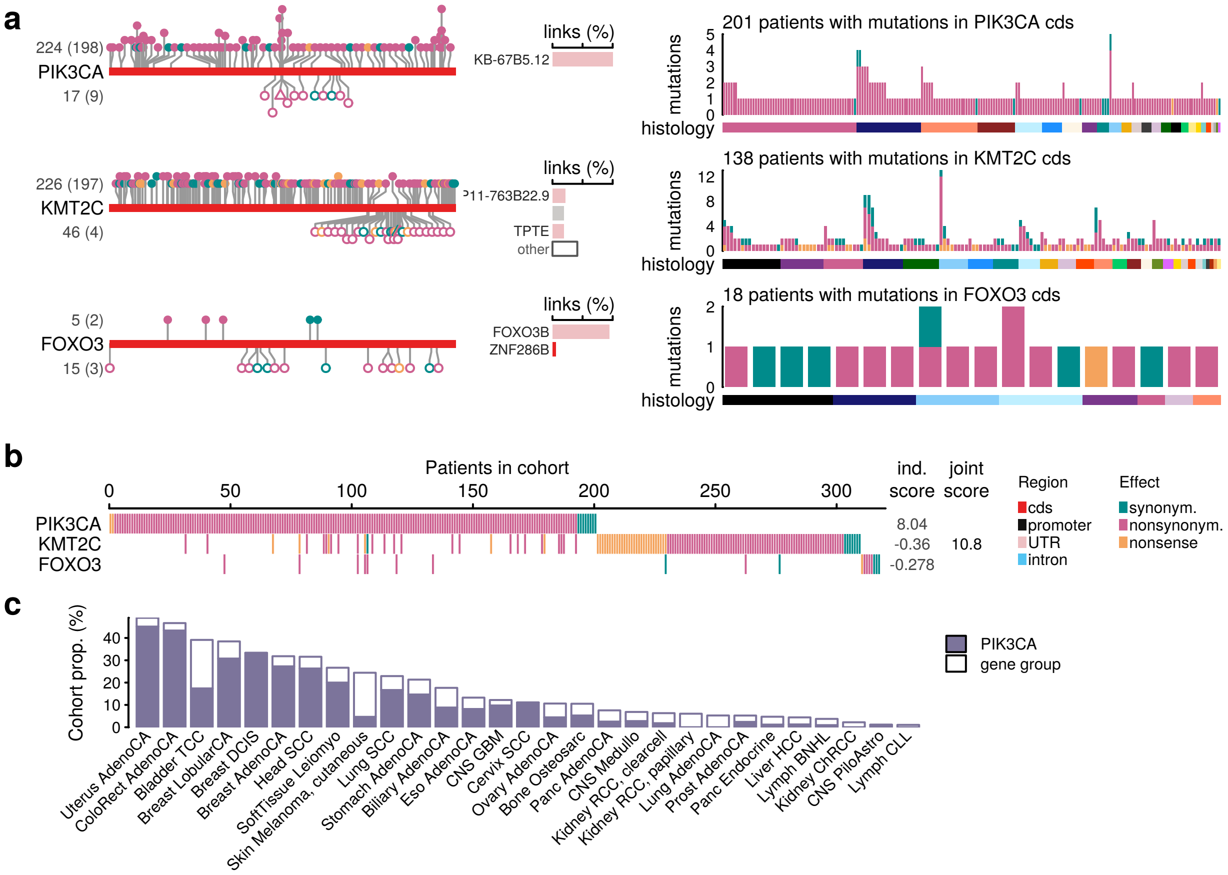


**Figure S14.** Cohort view of selected cancer genes. **(a)** Distributions of mutations along selected genes. Panels on the right reveal how the mutations are partitioned among patients and cancer types. **(b)** Cohort view of the gene group in (a). For patients with more than one mutation in a gene, only one the most severe consequence is shown (nonsense, nonsynonymous, synonymous). **(c)** Proportions of patients in each cancer type carrying at least one mutation in the leading gene and in the gene group. This summary includes all samples, including hyper-mutator samples.

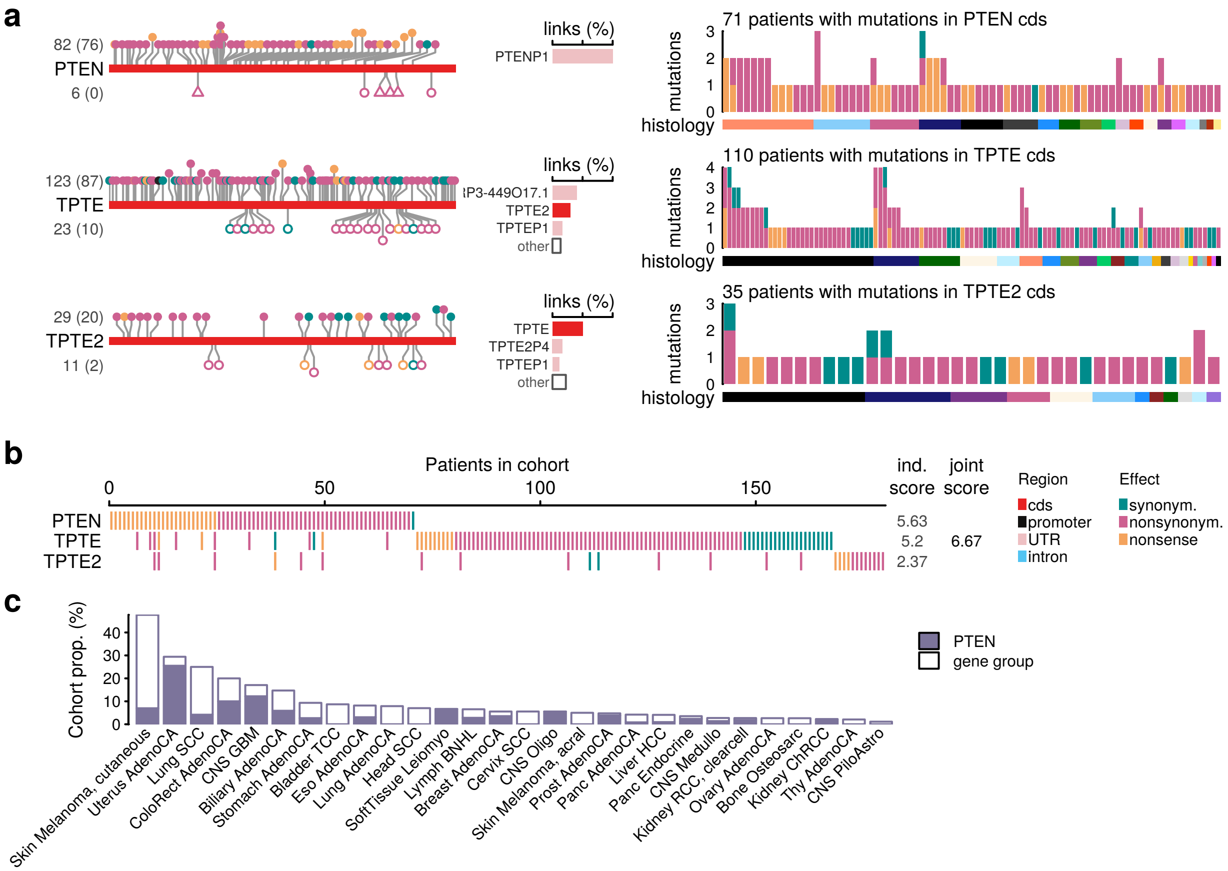


**Figure S15.** Cohort view of genes pertaining to the PTEN pathway.

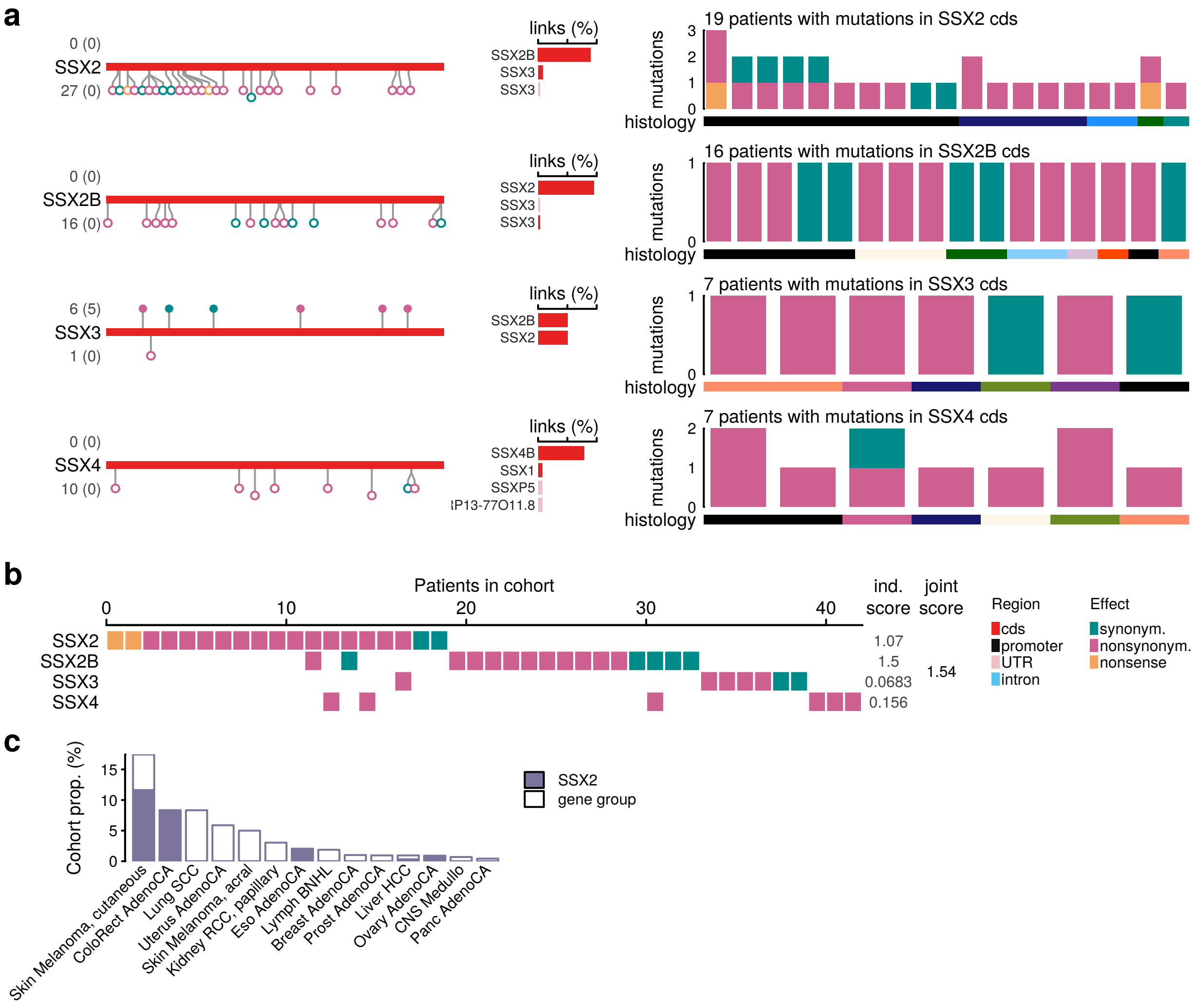


**Figure S16.** Cohort view of the SSX gene family.

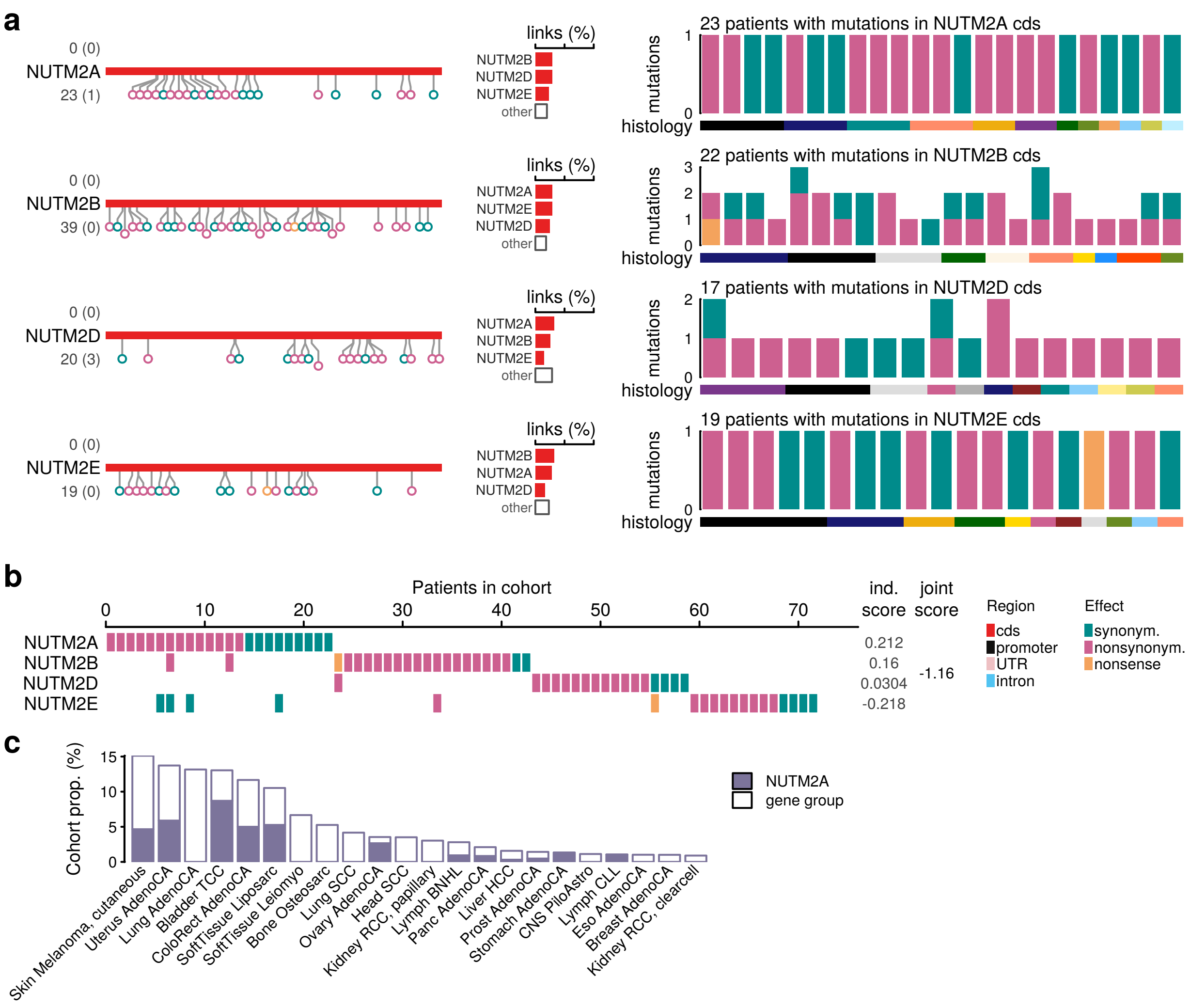


**Figure S17.** Cohort view of the NUTM2 gene family.

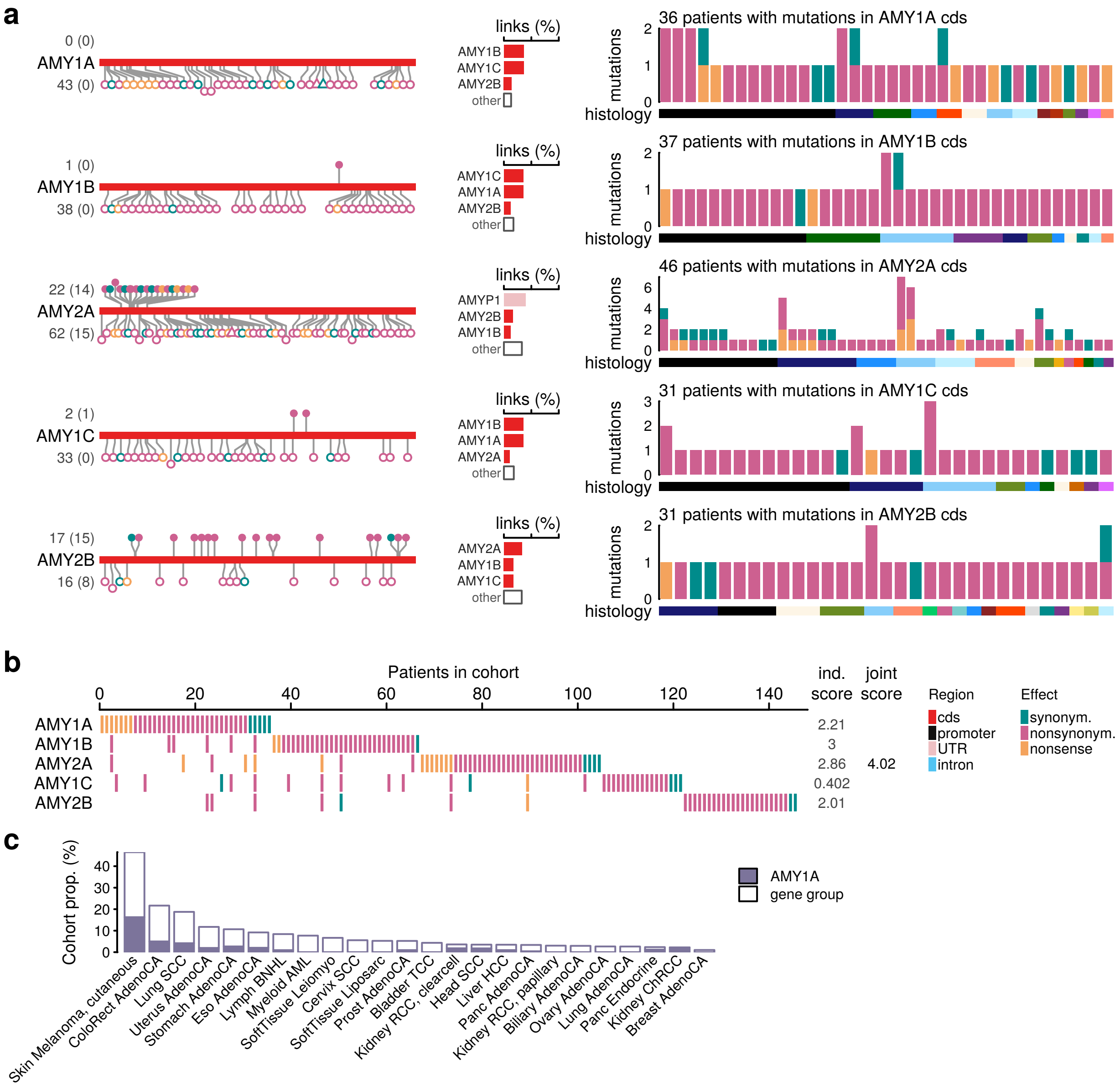


**Figure S18.** Cohort view of a group of genes from the AMY gene family.

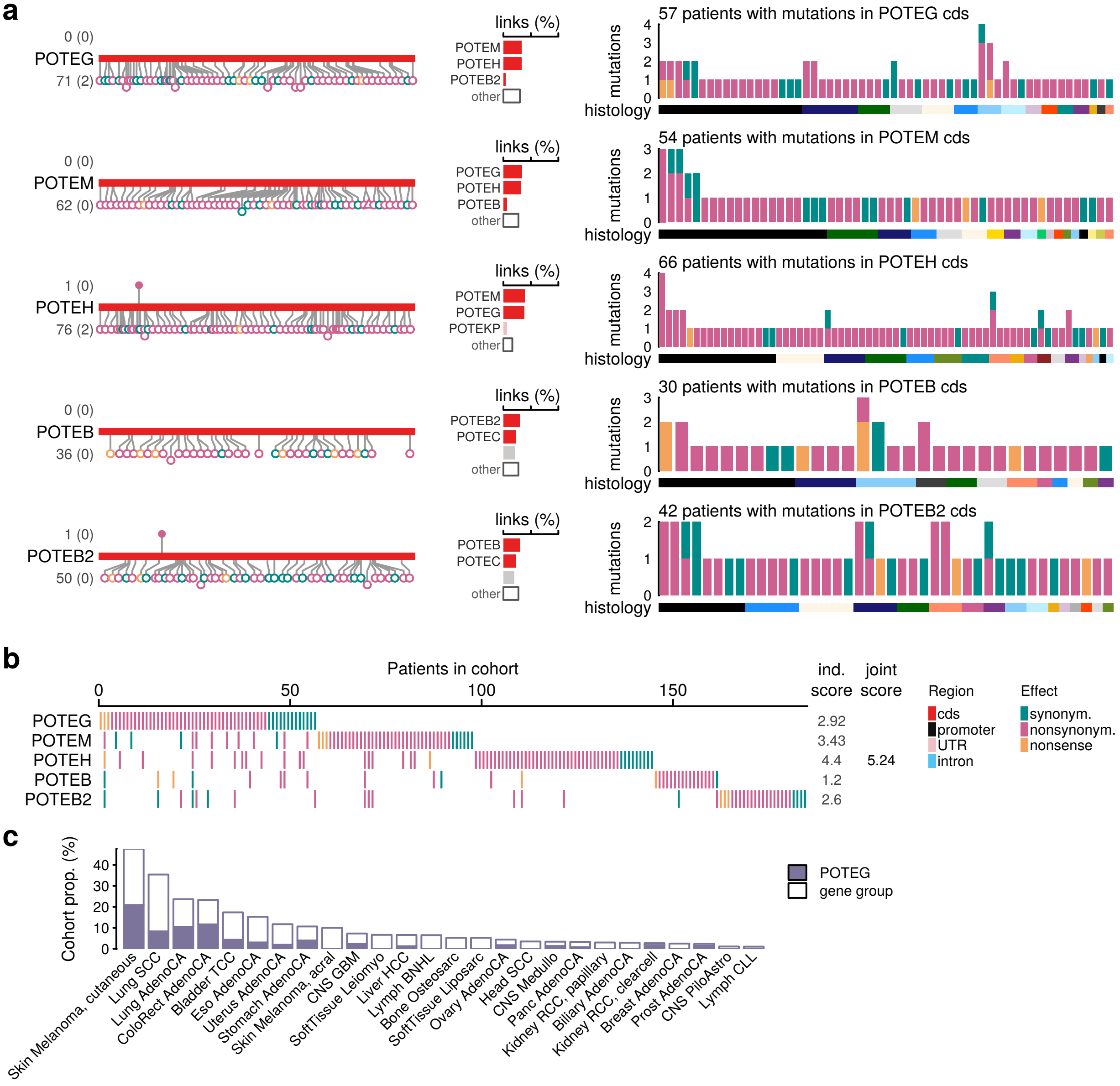


**Figure S19.** Cohort view of a group of genes from the POTE gene family.

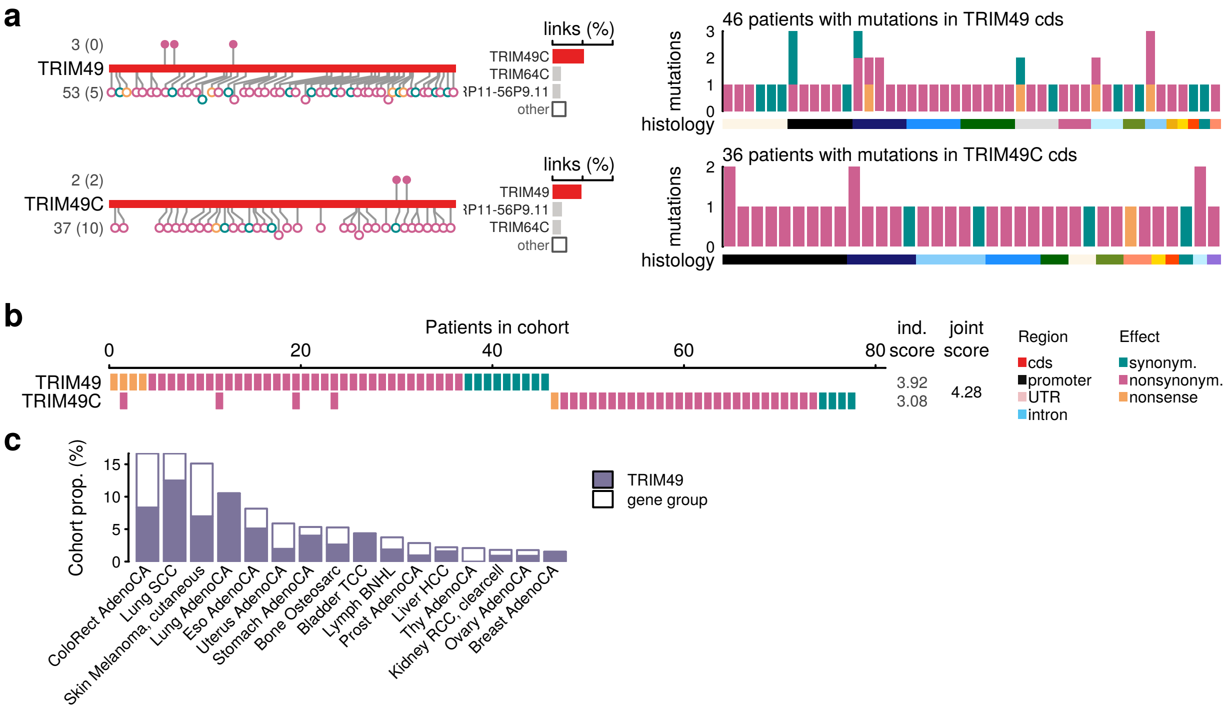


**Figure S20.** Cohort view of a group of genes from the TRIM gene family.

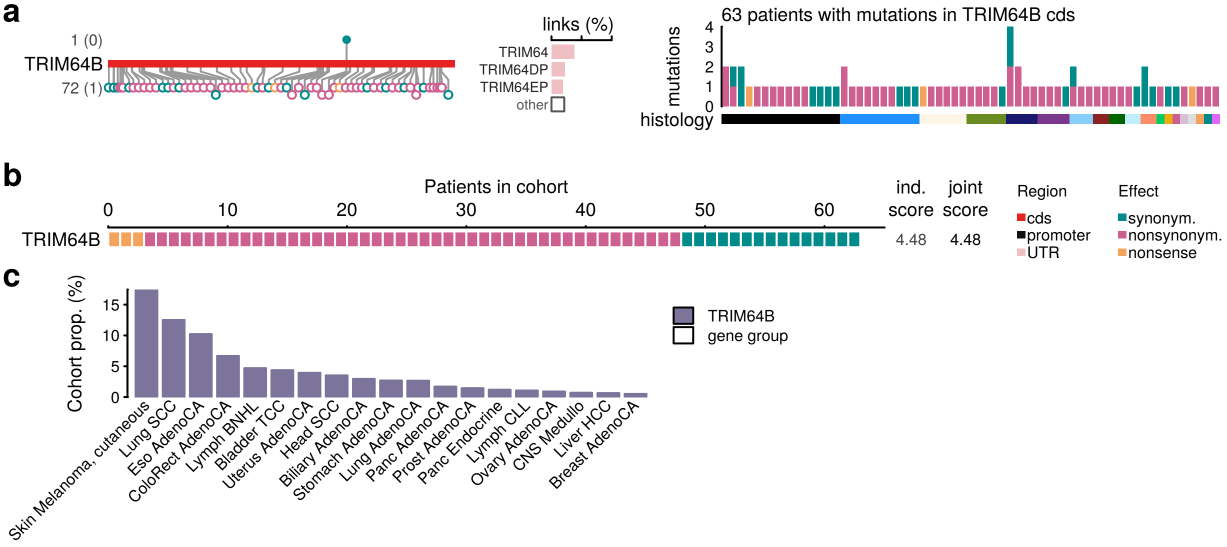


**Figure S21.** Cohort view of a genes in the TRIM gene family.

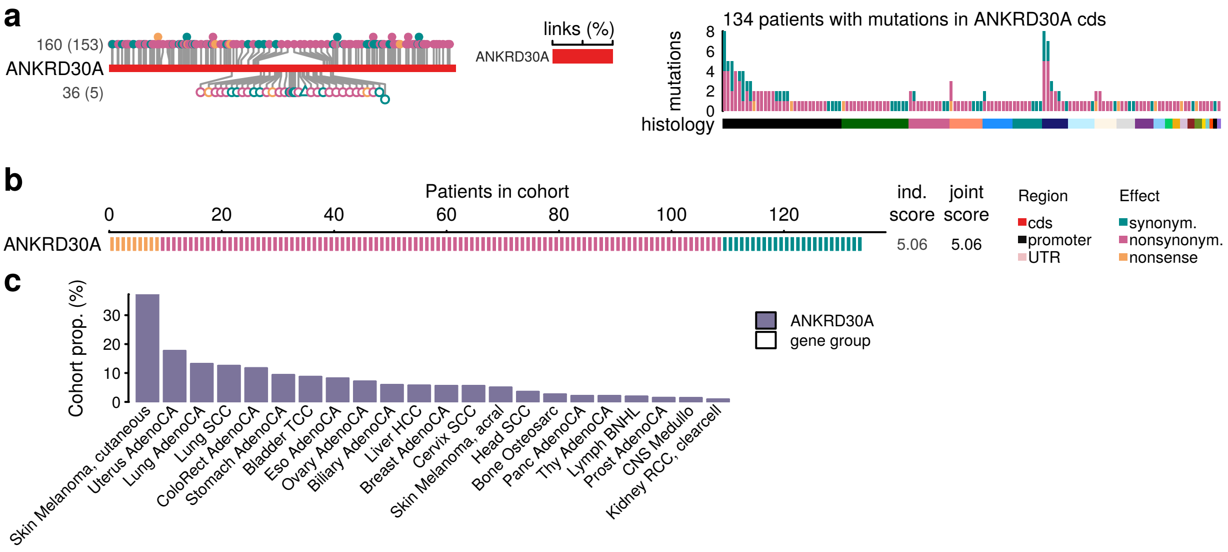


**Figure S22.** Cohort view of a gene in the ANKRD family.

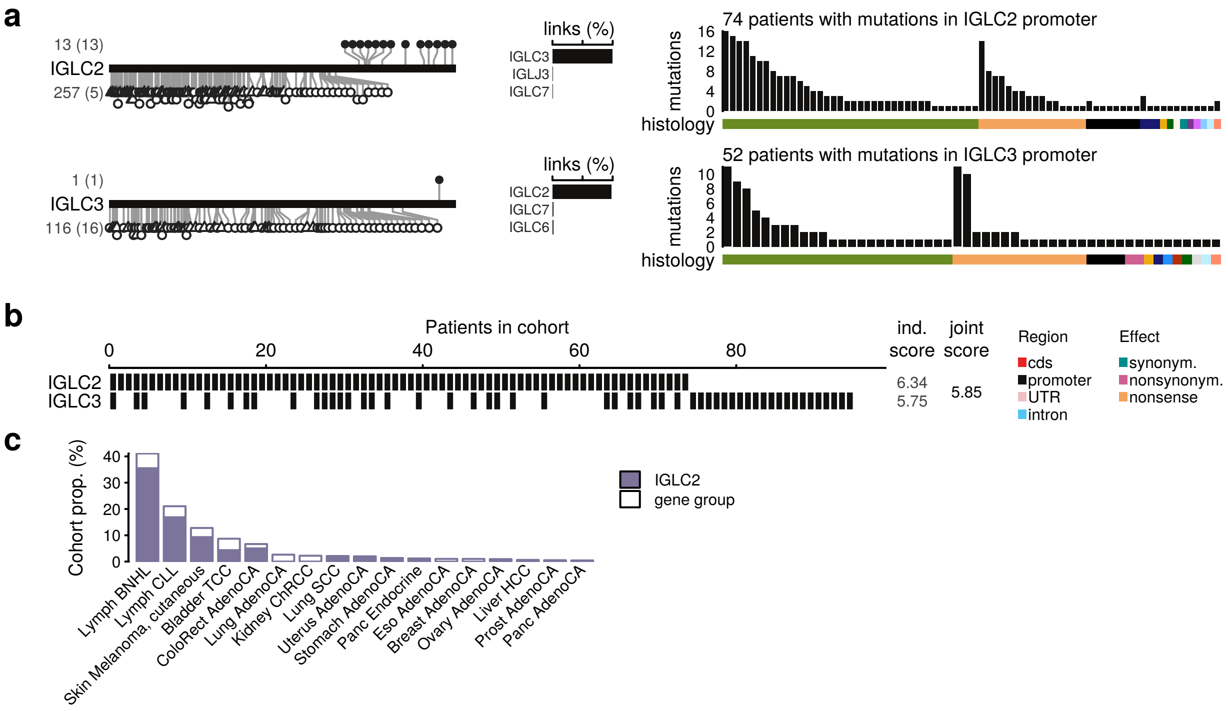


**Figure S23.** Cohort view of a group of promoters (upstream sequences) of the IGLC family.

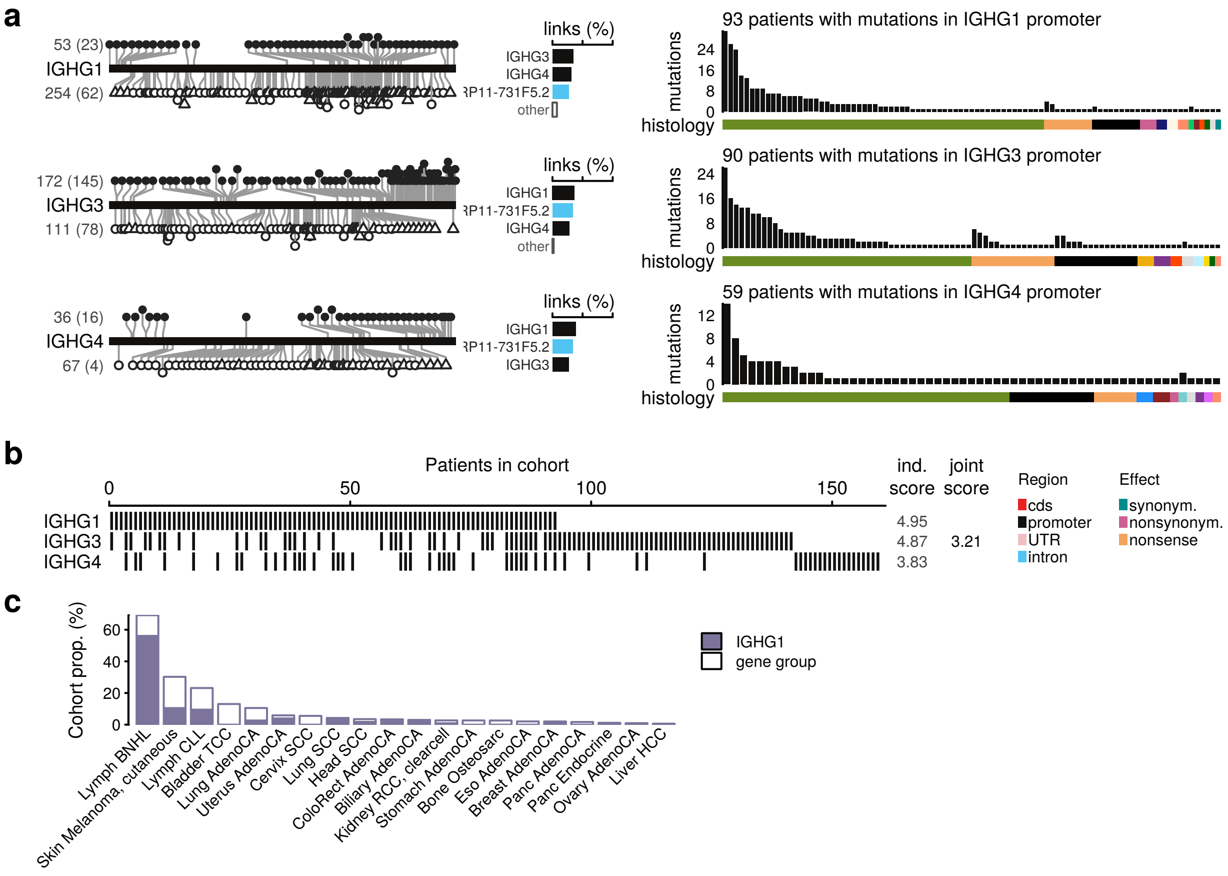


**Figure S24.** Cohort view of a group of promoters (upstream sequences) of the IGHG family.
